# Enhanced cell viscosity: a new phenotype associated with lamin A/C alterations

**DOI:** 10.1101/2022.07.18.500411

**Authors:** Cécile Jebane, Alice-Anaïs Varlet, Marc Karnat, Lucero M. Hernandez-Cedillo, Amélie Lecchi, Frédéric Bedu, Camille Desgrouas, Corinne Vigouroux, Marie-Christine Vantyghem, Annie Viallat, Jean-François Rupprecht, Emmanuèle Helfer, Catherine Badens

## Abstract

Lamin A/C is a well-established key contributor to nuclear stiffness and its role in nucleus mechanical properties has been extensively studied. However, its impact on whole cell mechanics has been poorly addressed, even less so in terms of measurable physical parameters. In the present study, microfluidic experiments combined with theoretical analyses were performed to provide a quantitative estimation of the whole cell mechanical properties. This allowed the characterization of mechanical cell changes induced by lamin A/C alterations resulting from Atazanavir treatment or lipodystrophy-associated LMNA R482W pathogenic variant. Results unveil an increase in the long-time viscosity as a signature of cells affected by lamin A/C alterations. In addition, they show that the whole cell response to mechanical stress is driven not only by the nucleus but also by the nucleo-cytoskeleton links and the microtubule network. This enhanced cell viscosity assessed by our microfluidic device could represent a useful diagnosis marker for lamin-related diseases.

## Introduction

Lamin A/C is an intermediate filament underlying the nuclear envelope and a well-established key contributor to nuclear stiffness, chromatin organization and gene expression (Isermann and Lammerding, 2013; Hah and Kim, 2019). Due to its links with both the nucleoskeleton and the cytoskeleton, lamin A/C is also a major piece of the mechanotransduction process, i.e. the conversion of mechanical stimuli into biochemical signals (Kalukula et al., 2022; Zuela-Sopilniak and Lammerding, 2022). With regard to these fundamental and ubiquitous roles, lamin A/C alterations resulting from genetic or environmental factors, induce diverse cellular features such as abnormal nuclear shape, proliferation defect and premature senescence (Dechat et al., 2008; Kim et al., 2019; Varlet et al., 2020). Genetic diseases due to pathogenic variants in LMNA, the gene encoding Lamin A/C, are referred to as laminopathies. They are severe and clinically heterogeneous diseases either tissue-specific like cardiomyopathies, or multi-systemic like Hutchinson-Gilford progeria syndrome and type 2 familial partial lipodystrophy (FPLD2) (Kang et al., 2018; Shin and Worman, 2022; Zammouri et al., 2022).

On account of their pivotal role in nuclear stiffness, the involvement of lamins in nucleus mechanical properties has been extensively studied through a variety of techniques applied to different cell types and different lamin modifications such as depletion or genetic variants. However, the diversity of approaches has resulted in a large panel of results that are rather difficult to compare (Crasto et al., 2020; Kalukula et al., 2022; Urciuoli and Peruzzi, 2022). In addition, the link between lamin A/C structural or functional changes and cell or tissue manifestations in the pathological context of laminopathies, remains unclear. To a large extent, the impact of laminopathies on the whole cell mechanical properties has been poorly described in terms of measurable physical mechanical parameters since most studies have focused on nucleus investigations (Isermann and Lammerding, 2013; Varlet et al., 2020).

Here, the mechanical impact of two types of lamin A/C alterations was investigated: (1) the canonical FPLD2-associated lamin A/C R482W pathogenic variant and (2) an *in vitro* treatment with Atazanavir, a HIV protease inhibitor known to induce senescence phenotypes associated to lamin A/C alterations (Caron et al., 2007; Bonello-Palot et al., 2014; López-Otín et al., 2013). To test the whole cell response to mechanical stress, the microfluidic technique, largely applied to infer cellular mechanical properties, was implemented (Davidson et al., 2015; Herbig et al., 2018; Dupire et al., 2020). A microfluidic device designed to apply a compressive stress on cells by forcing them into constrictions, was used to provide a quantitative data set on the cell mechanics. The mechanical stress was applied over a few seconds to probe the intrinsic cell properties without inducing a major reorganization of cellular components. Regarding this aspect, the study differs from the majority of previously reported experiments in which the mechanical stress is applied over longer times ranging from 10 seconds to a few minutes (Lange et al., 2015; Davidson et al., 2019; Wintner et al., 2020). Measurements of the cell transit time in constrictions were combined with a semi-automated image analysis routine and the application of a rheological model to extract the cells’ viscoelastic properties. The results showed that the main mechanical feature of cells displaying lamin A/C alterations, whether induced *in vitro* or by the R482W mutation, is an increase in the cell’s viscosity attributing to both nuclear and cytoskeleton networks. Moreover, a contribution of the microtubule network in this mechanical property was unraveled in cells with lamin A/C alterations.

## Results

### Atazanavir treatment drives a premature senescence phenotype in fibroblasts identical to that induced by lamin A/C R482W mutation

Two cell models characterized by lamin A/C alterations were selected to study cell viscoelastic property changes as compared to control cells: 1) skin fibroblasts issued from a healthy individual and treated with Atazanavir, and 2) skin fibroblasts issued from two patients with FPLD2 carrying the canonical lamin A/C R482W mutation (patients 8 and 12 from Vatier et al. (Vatier et al., 2016)).

The response of control fibroblasts to Atazanavir was checked by studying the nuclear shape, proliferation capacity and senescent markers. Cells were treated for 48 h with increasing concentrations of Atazanavir, or incubated only with its solvent, DMSO, as a control. High levels of nuclear aberrations were observed and were similar to those observed in cells from patients with FPLD2 or from a patient with Progeria carrying the LMNA G608G mutation (Fig. 1A-C). The impact of increasing Atazanavir concentration was also confirmed on proliferation capacity by measuring Bromodeoxyuridine incorporation, and on senescence level by estimating the cellular SA-beta-galactosidase activity (Fig. 1D-E). From these data, the concentration of 50 µM was determined to efficiently drive senescence while avoiding impaired cell viability.

**Figure 1.**
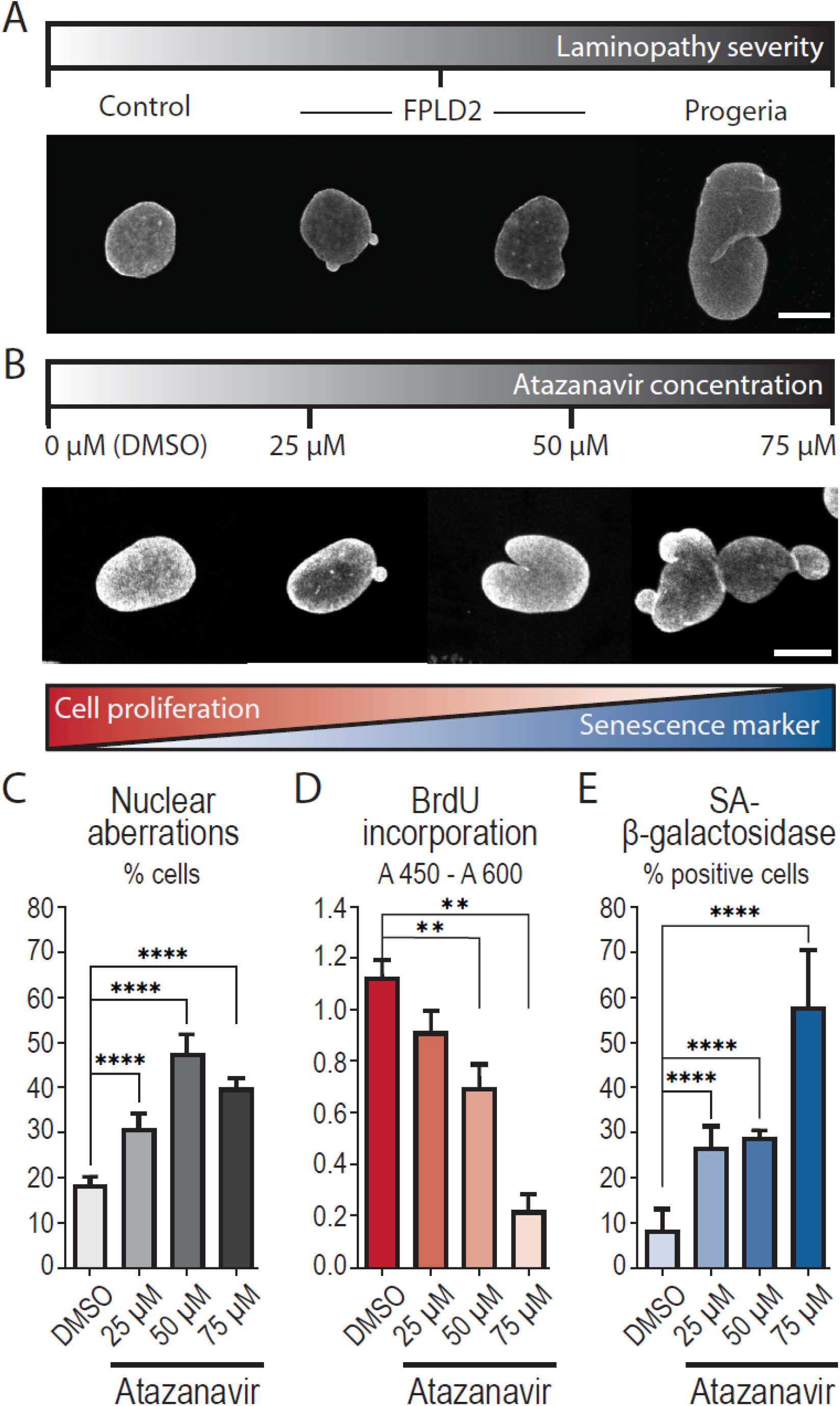
Atazanavir treatment induces cellular phenotypes associated with premature senescence. Control fibroblasts were treated for 48 h with increasing doses of Atazanavir or incubated with DMSO as a control. (A-B) Representative images of nuclear aberrations (abnormal size or shape) observed in (A) control fibroblasts, fibroblasts from patients with FPLD2 carrying the LMNA R482W mutation, or from a patient with Progeria, and (B) control fibroblasts treated with Atazanavir. Nuclear envelope is stained using a lamin A/C specific antibody. Scale bars: 10 µm. (C) Percentage of cells displaying nuclear aberrations. (D) Bromodeoxyuridine (BrdU) incorporation reporting cell proliferation ability. (E) Percentage of SA-beta-galactosidase positive cells identified as senescent. Errors are standard deviations. Number of experiments: N ≥ 3 (C-E); number of analyzed cells: n ≥1000 (C) and ≥ 500 (E).

As a second step, we evaluated the impact of Atazanavir treatment and FPLD2 mutation on cellular and nuclear volumes to engineer constriction dimensions in the microfluidic device described hereafter. Volumes were derived from confocal imaging of cells in suspension, to mimic the microfluidic condition. Cell and nuclei volumes of control fibroblasts treated with Atazanavir increased by 37% and 13%, respectively, when compared to those of fibroblasts incubated with DMSO only. The volumes of fibroblasts from the 2 patients affected with FPLD2 (referred to as K and M) were compared to those of control cells and to those of fibroblasts from a patient affected with Progeria, known to induce an increase in nuclear and cellular volumes (Goldman et al., 2004; Veitia, 2019). As expected, both cellular and nuclear volumes of Progeria cells were much larger than the others, whereas the volumes of FPLD2 cells and nuclei remained closer to those displayed by control cells (Fig. S1).

Altogether, our results suggest that Atazanavir treatment induces premature senescence in control fibroblasts, with similar nuclear dysmorphism as compared to cells bearing FPLD2-associated LMNA mutation. As their volumes are similar, cells treated with Atazanavir or carrying the FPLD2 mutation and control cells can be compared in the same microfluidic experiments, conversely to larger Progeria cells.

### A two-height microfluidic device to infer the cell responses to mechanical constraint

To investigate cell mechanical properties, we engineered a microfluidic device consisting of a wide channel with parallel constrictions dedicated to constraining cells (Fig. 2A-B). An original feature of the device lies in the presence of two heights along the channel (Fig. 2B): in the center of the device lies a region with constrictions of a 6×6 µm^2^ cross-section, which are small enough to squeeze both the cells and their nuclei. Before and after the central constrictions, the channel displays a 200-µm long section of a 6×300 µm^2^ cross-section. By applying a controlled pressure difference between inlet and outlet, cells are pushed through channels and deform progressively as they enter the constrictions. This device allows to observe the passage of tens of cells through the constrictions and, therefore, to quantitatively analyze the cell behavior. The dynamics of cell entry into the constrictions is measured by the temporal evolution of cell portion inside the constriction called cell tongue (Fig. 2C and Video S1). This dynamics exhibits three different successive phases, I to III (Fig. 2D) each being characterized by a specific linear increase in cell tongue length over time, i.e., a specific entry velocity. Once the cell has fully entered the constriction, its deformation is maximum and its transit through the constriction corresponds to phase IV (Fig. 2D).

**Figure 2.**
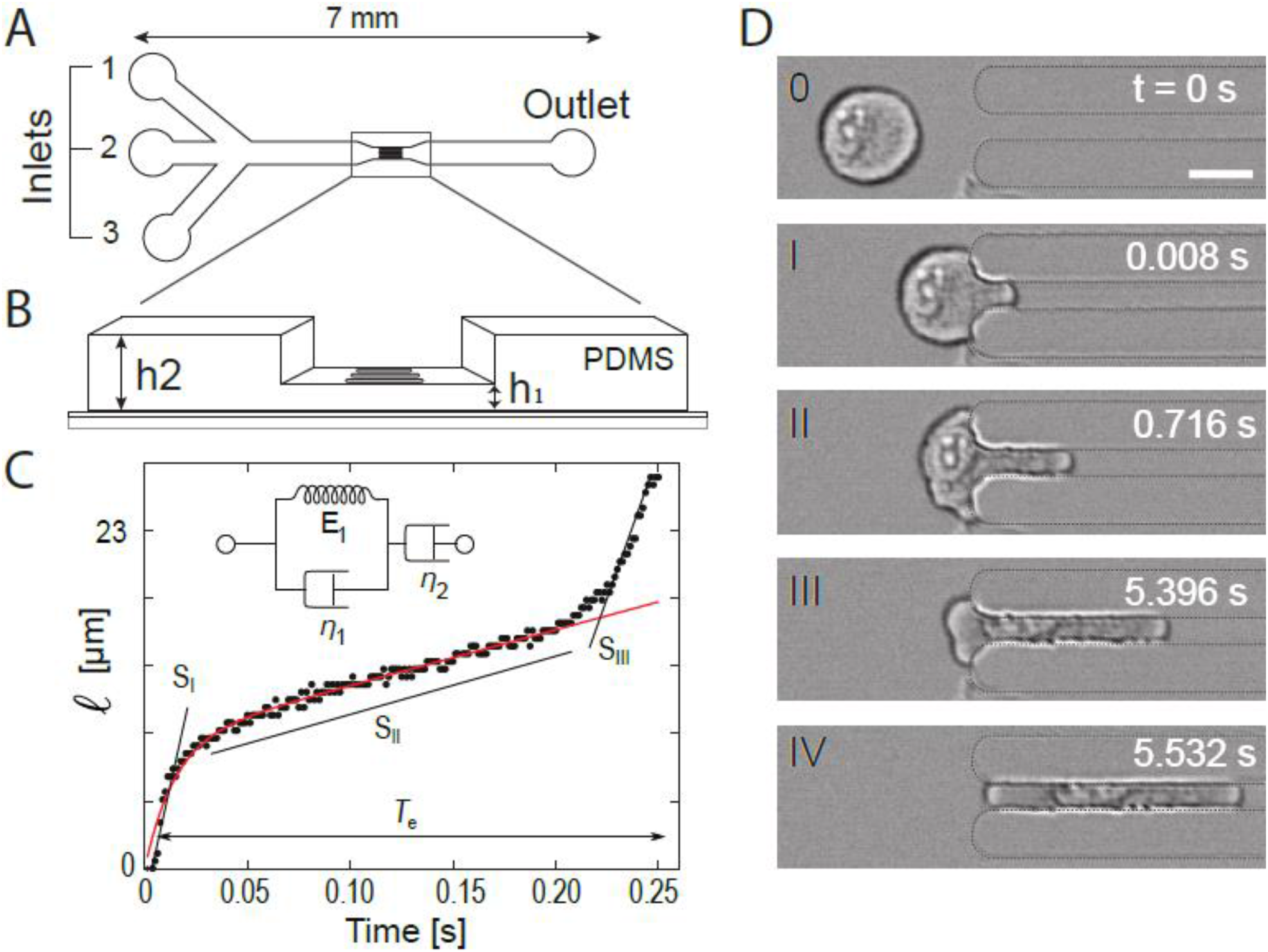
Microfluidic device, data processing and analysis. (A) Top-view schematics of the microfluidic device made of polydimethylsiloxane (PDMS). Inlets (1) and (3) are used for injecting buffer solutions while cells are injected through inlet (2). Controlled pressure drop is applied between inlets and outlet to push cells through constrictions. (B) Zoomed 3D-view of the constriction zone, with two different heights: h_2_ for the main 300-µm wide channel and h_1_ for the constrictions (length l, width w). l = 100 µm, w = h_1_ = 6 µm, h_2_ = 10 µm. The low-height region is 500 µm long in total, including the constrictions. (C) Typical time evolution of the cell tongue length l(t). The curve is analyzed using two methods: i) It is decomposed into three linear parts, of slopes S_I_ to S_III_, which correspond to the cell entry time T_e_; ii) The first two parts of the curve are fitted (in red) using a rheological model which combines a Kelvin–Voigt solid (a spring of constant E_1_ in parallel with a dashpot of viscosity η_1_) in series with a dashpot of viscosity η_2_ (see inset). (D) Timelapse of a 24-µm cell entering a constriction of width w. Scale bar: 15 µm.

### Cells exposed to Atazanavir treatment or carrying the lamin A/C R482W mutation display an increase in viscosity

Time course curves of cell tongue length were analyzed in two ways: first, by determining the global entry time T_e_, defined as the time to go from phase I to III, and the three cell entry velocities corresponding to the three slopes S_I_ to S_III_, of the three phases; second, using a rheological model to fit phases I and II. Fibroblasts from the 2 patients with FPLD2 (named M and K) were compared to untreated control fibroblasts (UNT) and fibroblasts treated with Atazanavir (AZN) to fibroblasts incubated with DMSO only (DMSO). As the major metabolic signs of FPLD2 are dyslipidemia and diabetes, we also used fibroblasts from a patient with dyslipidemia and diabetes but without lamin A/C alteration (patient 17 from Dutour et al. (Dutour et al., 2011), named T2D hereafter) to validate that the assay specifically discriminates cells with lamin A/C alterations.

The entry times (T_e_) of UNT and DMSO cells were similar (0.4 and 0.46 s), confirming that the DMSO used as Atazanavir solvent had no effect on T_e_. In contrast, the entry times of AZN, M and K cells were at least 3 times longer than those of their respective controls (Figs. 3A, S2A and Video S2). The entry time of T2D cells was slightly longer than that of UNT cells but significantly shorter than that of FPLD2 or AZN cells (Figs. 3A and S2A). To check that these variations in entry times were not due to a cell volume effect, cells were sorted by volume. As expected, entry times were longer for larger cell volumes, but for comparable volumes, those of FPLD2 and AZN cells were systematically longer than those of their respective controls (Fig. S2B).

**Figure 3.**
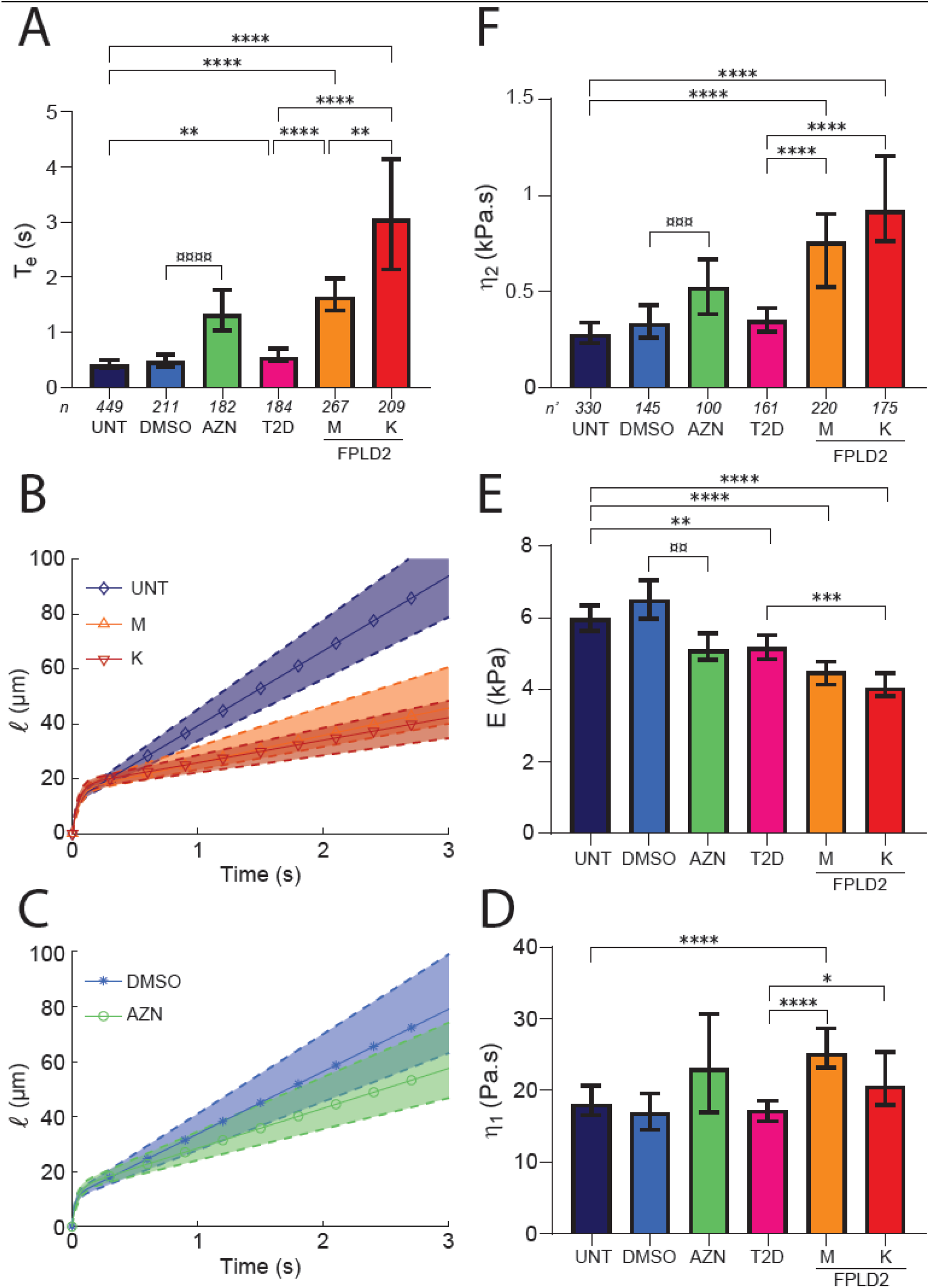
Atazanavir treatment and lamin A/C R482W mutation alter cell rheological behavior. UNT: untreated control cells (control); DMSO: control cells incubated with DMSO (AZN control); AZN: control cells treated with Atazanavir; T2D: cells from a patient with type 2 diabetes; M,K: cells from patients with FPLD2 carrying the LMNA R482W mutation. (A) Entry time in 6×6-µm^2^ constrictions T_e_ (median ± 95% CI). (B-C) Fits of tongue length l as a function of time t (t = 0 defined at cell contact with constriction), for (B) UNT, M and K cells, and (C) DMSO and AZN cells. Solid curves with symbols indicate the median fits; dashed upper and lower curves delineate the 95% CIs; the curves are plotted using median and CI values from Table 1 in Eq. (1). (D-F) Fit-extracted rheological parameters: long-time viscosity η_2_ (D), short-time elastic modulus E (E) and viscosity η_1_ (F). Medians ± 95% CI are displayed. Significant differences: AZN vs DMSO cells (¤), FPLD2 and T2D vs UNT cells (*). Number of experiments: N ≥ 3; n: number of analyzed cells (A); n’: number of fitted curves (D,E,F).

The cell entry velocities of the three phases (slopes S_I_ – S_III_) were slowed down for M and K cells compared to UNT cells, as well as for AZN cells compared to DMSO cells, in agreement with the observed increase in entry times (Fig. S3). For T2D cells, the entry velocities were similar to those of UNT cells in phase I and II, and higher in phase III, indicating again that their response to mechanical stress is different from that of FPLD2 and AZN cells. Altogether, these results showed that entry times, overall or for the different phases, were longer for cells with lamin A/C alterations, indicating changes in their mechanical properties.

To confirm this assumption and quantitatively measure the cell mechanical properties, the Jeffreys rheological model was used as a complementary approach ((Guevorkian et al., 2010), see Materials and Methods). This rheological model considers the cell as a homogeneous viscoelastic material with viscoelastic properties like elastic modulus and viscosities which respectively represent its ability to resist to deformation and its deformation speed. The model was used to fit the first 2 entry phases I and II only because the differences in entry phase III could not be clearly attributed to cell conditions. As the − minimal pressure difference required to push cells through the constrictions was estimated and found relatively minor compared to the pressure difference applied during experiments, the model was simplified by considering the applied stress as constant. Assuming constant applied stress on the cell and volume conservation, the cell deformation during constriction entering, i.e., the cell tongue length, can be expressed as a function of the cell viscoelastic properties using the Jeffreys model (Guevorkian et al., 2010; Davidson et al., 2019). The evolution of the cell tongue length *l* according to time *t* then reads as:

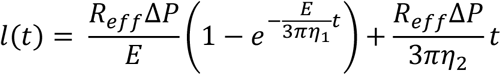

where R_eff_ is an effective cylindrical radius equivalent to the squared cross-section of the constriction,ΔP the applied pressure difference, E the elastic modulus of the cell, and η_1_ and η_2_ its short- and long-time viscosities. Following this model, at small deformations associated with short-time scale (phase I), a viscoelastic regime dominates which is controlled by the short-time viscosity η_1_ and the elastic modulus E. At larger deformations and long-time scale (phase II), a linear viscous regime dominates controlled by the long-time viscosity η_2_.

The cell tongue lengths obtained from experiments on individual cells were thus fitted using the Jeffreys model to derive their viscoelastic properties. Figure 3B-C displays the fits for UNT vs M and K cells, and for DMSO vs AZN cells, respectively, plotted using the median and CI values of the viscoelastic properties. In phase I, the short-time viscosity η_1_ tended to be higher in altered lamin A/C cells compared to their respective controls, and it was significantly different between FPLD2 cells and T2D cells (Fig. 3D). The elastic modulus E, on the other hand, was lower in AZN and FPLD2 cells, by 22 to 32% compared to their respective controls (Fig. 3E). T2D cells also displayed a slightly lower elastic modulus suggesting that the changes are not related to lamin A/C alterations only. At long-time scale, in phase II, UNT, DMSO and T2D cells presented a comparable long-time viscosity η_2_, while AZN and FPLD2 cells displayed an increase in η_2_, by at least 50% (Fig. 3F). Entry time T_e_ and viscosity η_2_ showed a similar trend with an increase in cells carrying altered lamin A/C, suggesting that the entry time is mostly dominated by the long-time viscosity. All rheological parameters are summarized with T_e_, in Table 1 (the corresponding dot plots are shown in Figs. S4 and S2A).

**Table 1.** Entry time and rheological parameters (Median [95% CI]). Colors indicate significant differences: AZN vs DMSO cells, ns|*|**|***|****; T2D/M/K vs UNT cells, ns|*|**|***|****.

|  | UNT | DMSO | AZN | T2D | M | K |
| --- | --- | --- | --- | --- | --- | --- |
| **T<sub>e</sub>**<br>**(s)** | 0.40<br>[0.35-0.50] | 0.46<br>[0.37-0.60] | 1.30<br>[1.00-1.77] | 0.54<br>[0.48-0.70] | 1.60<br>[1.40-1.98] | 3.05<br>[2.14-4.14] |
| **η<sub>1</sub>**<br>**(Pa.s)** | 18<br>[17-21] | 17<br>[15-20] | 23<br>[16.6-31] | 17<br>[16-19] | 25<br>[23-29] | 21<br>[18-25] |
| **E**<br>**(kPa)** | 6.0<br>[5.6-6.3] | 6.5<br>[6.0-7.0] | 5.1<br>[4.8-5.6] | 5.2<br>[4.8-5.5] | 4.5<br>[4.1-4.8] | 4.1<br>[3.8-4.5] |
| **η<sub>2</sub>**<br>**(Pa.s)** | 277<br>[232-336] | 333<br>[260-428] | 521<br>[381-668] | 349<br>[292-413] | 758<br>[522-904] | 920<br>[762-1206] |

With additional simplifications in the Jeffreys model (see Materials and Methods for details), we obtained a scaling law between the entry time and the cell initial diameter, T_e_ = ν·L0^2^, which could be used to fit the T_e_(L_0_) data sets and derive the factor ν (see, for example, the fits for UNT and K cells in Fig. S5A). The ν values normalized to that of UNT cells (ν/ν_UNT_) behave almost identically to the η_2_ ratios (η_2_/η_2_UNT_) obtained with the precise fits of the elongation curves (Fig. S5B). Thus, the measurement of the cell entry time and its initial size seems appropriate to perform a first rapid comparison of the long-time viscosities between different cell types. This method constitutes a fast screening of cell populations to obtain a rough estimation of viscosity differences based on the sole measurement of the entry time and initial cell diameter.

We also applied a power-law fit to the curves as in Davidson et al. (Davidson et al., 2019) which derives a parameter equivalent to the long time viscosity obtained with the Jeffreys model. This second approach provided results with a similar trend, though less marked (see Materials and Methods and Fig. S6).

Taken altogether, our data highlight that lamin A/C alterations, induced either by Atazanavir treatment or FPLD2-associated R482W mutation, are associated with a higher long-time cell viscosity. The short-time viscosity values, which are much lower than those of the long-time viscosity, are comparable to the viscosity values previously reported for cytoplasm (Moeendarbary et al., 2013; Wang et al., 2019). This short-time viscosity is probably due to the formation of a bleb at the cell front when it contacts the constriction, a process that is expected not to depend on the nucleus (Fig. S7). Importantly, using this approach we were able to discriminate T2D cells, from a patient with type 2 diabetes, from FPLD2 cells, supporting that the mechanical phenotypes observed in phases I and II are related to LMNA pathogenic variants.

So far, the observed behavior corresponded to the whole cell response, with the combined contributions of both the nucleus and the cytoplasmic content. The lamin A/C alterations induced changes not only in phase II which was presumably attributed to the deformation of the nucleus entering the constriction but also in phases I and III, associated with the entry of front and rear cytoplasm, suggesting that alterations in the nuclear envelope impact the response of the whole cell and not only of the nucleus. To determine which part could be attributed to changes in nucleus only, the intrinsic properties of isolated nuclei were investigated.

### The lamin A/C-dependent whole cell response to mechanical constraint does not depend exclusively on the nucleus

The same microfluidic experiments, albeit with 3×3 µm^2^ squared constrictions, were performed on nuclei extracted from UNT, DMSO, AZN and both FPLD2 cell lines. The nucleus tongue length was measured over time and the nucleus entry time T_e_ was extracted.

The entry of isolated nuclei in constrictions was much faster than that of whole cells, within a few hundreds of milliseconds, without distinct entry dynamics (Fig. 4, Video S3). In accordance with the trend observed with whole cells, Atazanavir treatment increased nucleus entry time by approximately 50% compared to incubation with DMSO, regardless of the nuclear volume (Fig. S8A). Conversely, entry times of M and K nuclei were shorter than that of UNT ones, especially in the largest volume range (Fig. S8B), Assuming the entry time of isolated nuclei is dominated by their viscosity, as observed in whole cells, the results show that isolated nuclei behavior did not reproduce the response of whole cells. We thus tested a possible role of the altered nucleus-cytoskeleton interplay in modifying the viscosity of whole cells in the context of lamin A/C alterations.

**Figure 4.**
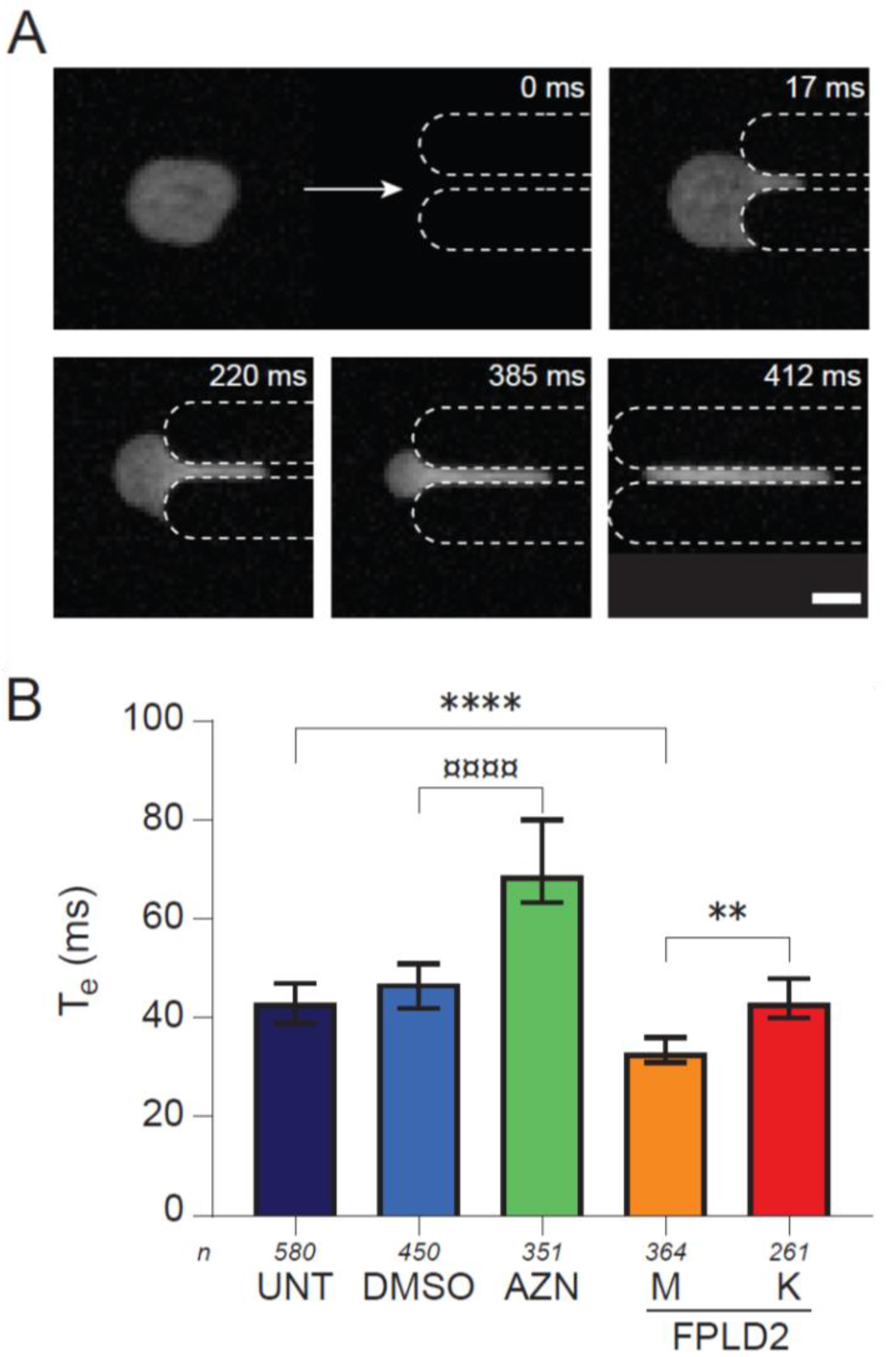
Nucleus response differs from whole cell response. (A) Timelapse of a 20-µm nucleus entering (I-III) and transiting (IV) through a 3×3-µm^2^ constriction. Nucleus is labeled with Hoechst. Scale bar: 10 µm. (B) Entry time of isolated nuclei in constrictions T_e_ (median ± 95% CI). Nuclei from UNT: untreated control cells (control); DMSO: control cells incubated with DMSO; AZN: control cells treated with Atazanavir; M,K: cells from patients with FPLD2. Number of experiments: N ≥ 4; n: number of analyzed isolated nuclei.

### Increase of cell viscosity in response to lamin A/C alterations implies the microtubule network

As actin and, to a lesser extent, microtubule networks are known to participate in cell viscoelastic properties, their contribution to cell responses was assessed by destabilizing actin and/or microtubule networks, using respectively latrunculin A or nocodazole treatment on UNT, AZN and FPLD2 K cells (Figs. 5A-B and S9). Just as for cells with an intact cytoskeleton, three phases were distinguished and the entry time T_e_ and viscoelastic parameters (E, η_1_, and η_2_) were quantified (Fig. 5C-N and Table S1). As expected, cytoskeletal drugs induced significant changes in cell response. Globally, the destabilization of either microtubule or actin networks induced a large decrease in T_e_, which remained higher for AZN and K cells than for control cells (Fig. 5C-E). Regarding the viscoelastic properties extracted by the rheological model, actin cytoskeleton disruption induced a significant decrease, by 40 to 65%, in both viscosities η_1_ and η_2_ in all three cell conditions and had a limited effect on the elastic modulus E in all studied cells, suggesting a similar role of actin (Fig. 5F-N).

**Figure 5.**
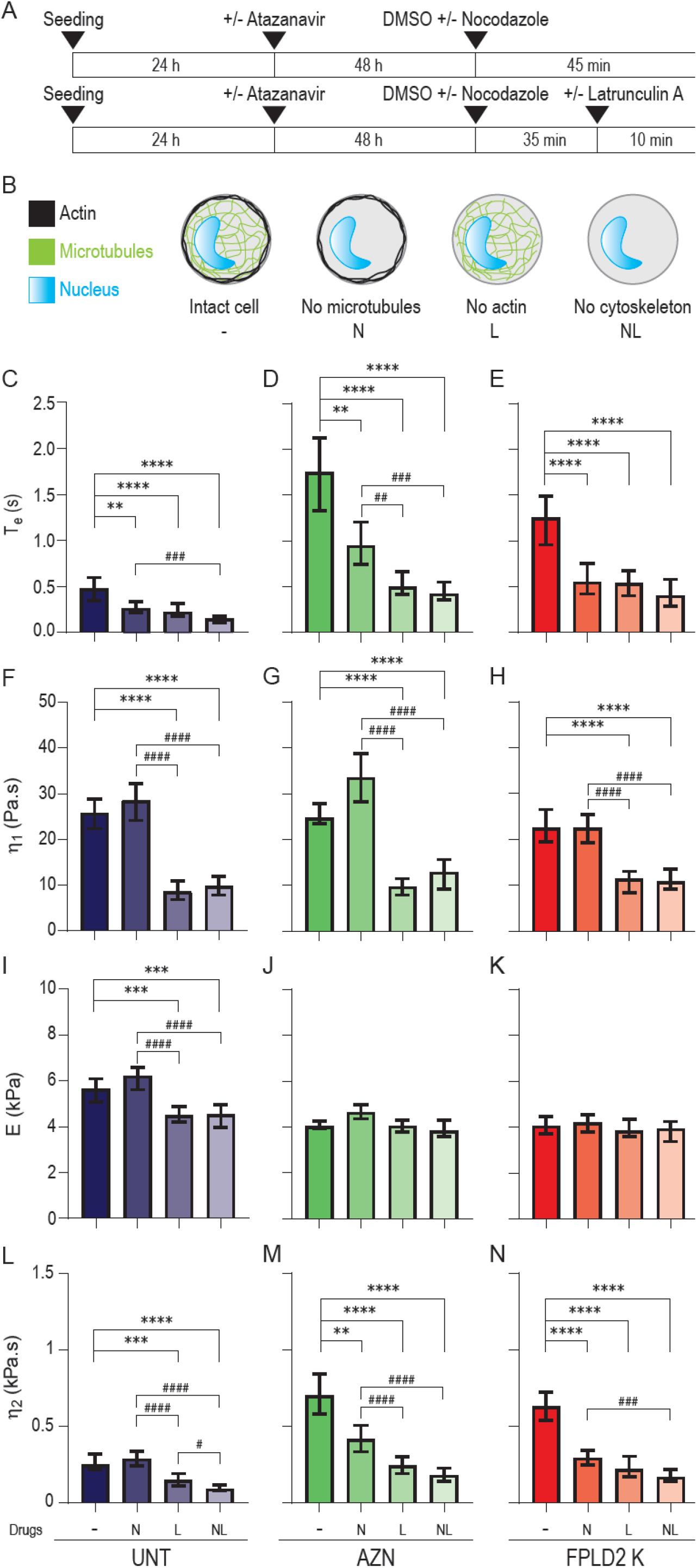
Actin and microtubule network destabilization accelerates cell entry into constrictions and decreases long-time viscosity in senescent cells. (A) Timeline protocol for cytoskeletal drug treatments: 72 h after seeding, cells, treated or not with Atazanavir, are treated with nocodazole (45 min), latrunculin A (10 min), or both (35 min with nocodazole, then 10 min with both drugs). (B) Sketched effect of drug treatments on cells, with nocodazole and/or latrunculin A leading to microtubule and/or actin destabilization. Control condition is incubation with drug solvent, DMSO. (C-E) Entry time in constrictions T_e_ and F-N) fit-extracted rheological parameters of untreated control cells (UNT, dark blue), Atazanavir-treated cells (AZN, green), and FPLD2 cells (FPLD2 K, red) upon drug treatments. Medians ± 95% CIs are displayed; significant differences: drug treatments vs DMSO control (*), and between drug treatments (^#^). Number of experiments: N ≥ 3; number of analyzed cells: n ≥ 100 (C-E); number of fitted curves: n’ ≥ 60 (F-N).

In response to microtubule network disruption only, short-time viscosity η_1_ and the elastic modulus E were barely impacted regardless of the cell conditions (Fig. 5F-K). Conversely, the long-time viscosity η_2_ was unchanged in UNT cells but significantly decreased by 50% or more in both AZN and K cells in response to microtubule destabilization (Fig. 5L-N). Therefore, this observation unravels an unexpected contribution of microtubules in the cellular viscosity of cells with lamin A/C alterations.

Finally, the effect of actin filaments and microtubules appears to be cumulative as destabilizing both actin and microtubule networks induced an approximately 70% decrease in entry time and long-time viscosity, regardless of cellular lamin A/C alterations. Altogether, our results suggest that the nucleus alone contributes, at most, by 30% to the viscous response of the whole cell, and that the effect of altered lamin A/C on cell viscosity mainly relies on cytoskeleton modifications.

## Discussion

In the present study, microfluidic experiments combined with a theoretical analysis were performed to explore changes in whole cell mechanical properties related to lamin A/C alterations caused either by Atazanavir treatment or FPLD2-associated LMNA R482W mutation.

It is largely assumed that alterations of the lamin A/C protein impact the mechanical properties of the nucleus. As such, previous studies have focused on cell nuclei only and used viscoelastic fluid models to report changes in nuclear viscosity and elastic modulus. For instance, the decrease of both nuclear viscosity and elastic modulus have been shown in lamin A/C depleted systems and in dilated cardiomyopathy-associated K219T mutation models (Davidson et al., 2019; Wintner et al., 2020). Upon lamin A/C overexpression or in Progeria-affected cells, an increased nuclear stiffness, a feature which can be interpreted as a higher elastic modulus, was reported (Dahl et al., 2006; Booth et al., 2015). In these studies, besides the fact that only nucleus changes were studied and that lamin A/C alterations were of different types, the observed timescale ranged from a few tens of seconds to a few minutes and the imposed deformation was usually limited to a few percent. In contrast, in our study, the dynamic response of the whole cells was assessed within at most a few seconds, and for high stresses and deformations (16.5 kPa pressure drop, 6×6 µm^2^ confinement). This high applied constraint does not aim to mimic physiological conditions such as deformations encountered in developing tissues or during cell migration like the migration of invasive cancer cells, but to provide measurable responses allowing the quantification of differences in cell mechanical properties, according to nuclear cytoskeleton composition. A relatively similar microfluidic assay was used by Lange et al., to derive what they called cell elasticity and fluidity from entry times in microconstrictions. They described increased cell elasticity and decreased cell fluidity upon overexpression of wild-type lamin A, features which could correspond to increased elastic modulus and viscosity using our analysis method (Lange et al., 2017).

Here, we showed that, in addition to the cell entry time, all three dynamic regimes of cell deformation were changed by lamin A/C alterations, with a significant slow-down of the second regime corresponding to an increase in cell viscosity. The changes in the viscous regime were not entirely reproduced with isolated nuclei, suggesting that the nucleus alone is not responsible for all changes observed in whole cells. Our results evaluate to 30%, at most, the nuclear contribution to the viscous response of the whole cell, and indicate that the cytoskeleton and its link with the nucleus play a major role in lamin A/C-dependent cell mechanical properties. The physical coupling between the cytoskeleton and the nuclear envelope is known to involve the Linker of Nucleoskeleton and Cytoskeleton (LINC) complex and its connections to lamin A/C through KASH domain proteins (Crisp et al., 2006; Brayson and Shanahan, 2017). However, this coupling can also result from direct interactions between cytoskeleton components and nuclear envelope proteins, such as microtubules which are linked to emerin (Salpingidou et al., 2007). The strong increase of the cell viscosity in lamin A/C altered cells could be due to the destabilization of the nucleus-actin coupling resulting from lamin A/C defects, and subsequent release of actin filaments normally bound to the nuclear envelope. Indeed, the bulk viscosity directly relates to the concentration of cytoplasmic actin filaments, as extensively shown by *in vitro* rheological studies (Janmey et al., 1991). In parallel, our experiments point out that the viscosity of cells with lamin A/C alterations is significantly decreased upon disruption of the microtubule network, which is not the case in control cells as expected the literature (Haga et al., 2000; Rotsch and Radmacher, 2000). Thus, microtubules, conversely to actin filaments, might remain anchored properly to the nuclear envelope when lamin A/C is altered, and even in a stronger way.

The data presented here show that the actin network is involved in short- and long-time viscosities, whereas it has only a weak contribution to elastic modulus changes, both in control cells and in cells with lamin A/C alterations. This suggests that the elastic modulus measured with our device might relate to additional cell components such as the intermediate filament vimentin known to regulate cell mechanical properties (Mendez et al., 2014; Lavenus et al., 2020; Pogoda et al., 2022)

In the Jeffreys model, the cell is considered as a homogeneous material whereas the nucleus is known to be stiffer than the cytoplasm (Guilak et al., 2000). Nevertheless, this global approach of the cell is largely used and allowed us to investigate the mechanical properties of the whole structure constituted by the cytoplasm, the nucleus and their physical links. Here, the model was used to fit the entry time of cells and extract rheological properties. Using this approach, the long-time viscosity η_2_ measurements were situated around 1 kPa and the elastic modulus E displayed values around 5 kPa, both measurements being consistent with the range of values reported in the literature (Mahaffy et al., 2004; Pogoda et al., 2012; Dulińska-Molak et al., 2014; Davidson et al., 2019; Sliogeryte and Gavara, 2019). Overall, the microfluidic test described here provides a quantitative estimation of the whole cell mechanical properties and unveils a specific mechanical signature of cells affected by lamin A/C alterations, namely an increase of the long-time viscosity and a critical contribution of the microtubule network in the cell mechanical properties. Moreover, for the first time, we have provided data on changes in cell mechanical properties resulting from the canonical lamin A/C mutation responsible for the most frequent type of genetic lipodystrophy, whereas, up to now, most studies on laminopathies have focused on LMNA pathogenic variants involved in cardiomyopathy, myopathy or Progeria.

Our study could have several implications for the management of human diseases. First, by showing the uncovered role of microtubules in the regulation of cell mechanics upon lamin A/C alterations, our results point to new fields of investigations regarding the pathophysiology of laminopathies. Second, given that altered deformability and changes in mechanical properties are key determinants in cell pathophysiology, as previously shown in several situations including invasive cancers, our device constitutes a step towards potential new inexpensive and standardized biomarkers to be transferred into clinical practice. In particular, although further investigations are required in cells with different LMNA pathogenic variants, enhanced cell viscosity assessed by a microfluidic device could represent a useful diagnosis marker for laminopathic diseases.

## Materials and Methods

### Cells, nuclei, treatments and senescence assays

#### Cells

Fibroblasts were issued from two patients affected with FPLD2 due to the c.1444C>T ; R482W heterozygous pathogenic variant of the *LMNA* gene (patients 8 and 12 in Vatier et al. (Vatier et al., 2016), referred to as patients K and M, respectively), and from a diabetic patient with no lamin A/C alteration (patient 17 in Dutour et al. (Dutour et al., 2011), referred to as patient T2D). Control and Progeria (HGPS) fibroblasts were purchased from Coriell Institute (AG07095, AG06917). Cell lines are summarized in Table S2. Cells were cultured at 37°C under 5% CO_2_ in low glucose Dulbecco Medium Eagle Modified (DMEM) supplemented with 15% fetal bovine serum (FBS) and 2 mM L-Glutamine (DMEM, FBS, L-Glutamine, Gibco), and used between passage 10 and 15.

#### Isolated nuclei

Nuclei were isolated following an adapted protocol from Kidiyoor et al. (Kidiyoor et al., 2020). Adherent cells were incubated in H_2_O MQ + IGEPAL 0.05% + Citric Acid 1% for 5 min at 37°C. The flask was vigorously tapped to expel the nuclei. The flask content was harvested, and the nuclei were washed with 10 mL PBS, vortexed ∼10 s and centrifuged at 800 g for 5 min. The supernatant was removed and the nuclei in the pellet were resuspended in PBS + BSA 1%. Nuclei were extracted right before microfluidic experiments.

#### Cell treatments

Atazanavir (stock solution at 50 mM in DMSO) was added at varying concentrations from 25 to 75 μM to the cell medium of control cells for 48 h before analyzing senescence-associated phenotypes. Control cells were incubated with 50 µM Atazanavir when performing microfluidic experiments. To induce the depolymerization of actin and/or microtubule networks, fibroblasts were treated with latrunculin A (stock solution at 1 mM in DMSO; L5163, Sigma-Aldrich) or nocodazole (stock solution at 50 mM in DMSO; 487928, Sigma-Aldrich). Cells were exposed to 200nM latrunculin A for 10 min, or 10 μM nocodazole for 45 min, or to both drugs (35 min nocodazole, then 10 min with latrunculin A and nocodazole), before being processed. For latrunculin A treatment, cells were incubated 35 min with DMSO prior to drug addition to keep 45 min total incubation with the carrier.

#### Senescence tests

Incorporation of 5-bromo-2’-deoxyuridine (BrdU): Cells were seeded on 96-well plate at a density of 10^4^ cells/well and BrdU was added to the cell culture medium 24 h before performing an ELISA assay following the manufacturer’s instructions (Cell proliferation ELISA, BrdU (colorimetric) Kit, Roche Applied Science). SA-β-galactosidase activity: Cells were seeded on glass coverslips (Lab-tek, SPL Life Sciences) coated with 100 μg/mL fibronectin (Sigma-Aldrich) and β-galactosidase activity was measured following the manufacturer’s instructions (Senescence β-Galactosidase Staining Kit, Cell Signaling Technology).

### Fluorescence microscopy, cellular and nuclear volume quantification

#### Immunofluorescence

Immunofluorescence was performed on adhered cells after fixation with 4% PFA at room temperature (RT) for 10 min and permeabilization with 0.5% Triton® X-100 (Sigma-Aldrich) at RT for 10 min. After washing twice with PBS, saturation was performed with 1% BSA (37525, Thermofisher) for 30 min at RT. Cells were incubated with 150 ng/mL Hoechst 33342 (H3570, Thermofisher) for 15 min at RT prior to saturation with primary mouse anti-lamin A/C (1:1000, sc-376248, Santa Cruz) or anti-α-tubulin (1:1000, T5168, Sigma-Aldrich) antibodies for 1.5 h at 37°C. After washing twice with PBS 0.1% tween, samples were incubated with an appropriate secondary antibody coupled with Alexa Fluor 488 (Invitrogen) for 1 h at 37°C. When required, actin staining was performed at the same time using Texas Red™-X Phalloidin (1:100, T7471, Thermofisher). Samples were then washed twice with PBS and post-fixed for 10 min with 4% PFA, before mounting on slides with ProLong™ Diamond Antifade Mountant (Thermofisher).

#### Nuclear aberrations

Immunofluorescence staining of lamin A/C was performed on fixed adhered cells and nuclear aberrations were quantified as previously described (Desgrouas et al., 2020). Nuclear phenotypes were monitored using an ApoTome system (Zeiss), equipped with a 100× objective and a CCD camera (Axiocam MRm, Zeiss). Nuclear abnormality criteria were aberrant nuclear lamin A/C staining pattern, enlarged nucleus, and aberrant nucleus shape. At least 1000 cells were examined for each condition and the percentage of abnormal nuclei was calculated.

#### Cellular and nuclear volumes

Adhered cells were labeled for plasma membrane and DNA by incubation for 1 h with CellBrite™ Green (Biotum) and 5 mg/mL Hoechst (Invitrogen), respectively, then detached using 0.05% Trypsin-EDTA (25300-054, Gibco). Cells were resuspended in complete cell medium supplemented with 1 mM HEPES and deposited on glass coverslips coated with 0.1 mg/mL PLL-PEG (PLL(20)-g[3.5]PEG(5), SuSoS AG) in 10 mM HEPES pH 7.4 to prevent cell adhesion. Cells in suspension were subsequently observed on a confocal microscope (LSM 800 airyscan Axio Observer Z1 7, Zeiss) equipped with a 63× water objective (63x/1.20 W Korr UV VIS IR C-Apochromat, Zeiss). Z-stacks were acquired with a 0.5-μm z-step in fluorescence channels corresponding to Cell Brite and Hoechst labeling. FIJI software (ImageJ) was used for volume quantification using a home-made macro: 1) the z-stack was thresholded; 2) for each stack slice *i* the object (cell or nucleus) contour was determined and its area Area_i_ was measured (in μm^2^); 3) the volume was calculated by summing the slice volumes: V = Σ_i_Area_i_·δh, with δh = 0.5 μm (z-step between slices). Nuclei are wrinkled when cells are in suspension, as previously reported (Kim et al., 2015; Lomakin et al., 2020). Here, the wrinkles were neglected in the estimation of the nuclear volumes.

### Microfluidic device fabrication

The microfluidic device is made of polydimethylsiloxane (PDMS, Dow Corning) and consists in a chamber assembled from a bottom PDMS-coated glass coverslip (defined as “bottom coverslip”) and a few mm-thick PDMS piece with drilled inlets/outlet (defined as “microchannel”).

#### Microchannel

The microchannel was fabricated from a master mold created on a silicon wafer using standard photolithography performed at PLANETE microfabrication facility of the laboratory (Fig. S11). Three masks were used to create the mold: (1) alignment crosses, (2) main channel with micrometer-sized constrictions in the central part, and (3) main channel without constrictions. A negative tone resist (AR-N 4340, AllResist) was spin-coated at 4000 rpm for 1 min on the silicon wafer before a first soft baking step at 85°C for 2 min, followed by exposure to UV for 14 s using mask (1) positioned with an aligner (MJB4 aligner, Carl Suss). A post-exposure baking step was performed (95°C, 2 min) before the resist layer was developed with a developer (AR300-475, Allresist). Next, a thin 160-nm aluminum (Al) layer was deposited by sputtering. A lift-off step was performed overnight with a solvent (NMP, MicroChemicals) to remove the resist which had not been exposed to UV and the Al deposited on it, leaving only the alignment crosses. To enhance adhesion between the resist and the silicon wafer, the wafer was plasma-treated for 10 min before an adhesion promotion layer (Omnicoat™, MicroChemicals) was spin-coated (3000 rpm, 40 s) then baked (200°C, 1 min). Permanent Epoxy Negative Photoresist (SU8-2005, MicroChemicals) was spin-coated on the wafer (1500 rpm, 60 s) to obtain a 6-μm layer before soft baking (9°C, 2 min). The resist was then aligned with mask (2) using Al crosses and exposed to UV for 10 s. The resist-coated wafer was then post baked (95°C, 3 min), developed in SU8-Developer (DevSU8/4, MicroChemicals) to remove unpolymerized resist, washed abundantly with isopropanol, air-blown and hard baked (cured) at 150◦C for 5-10 min to prevent the resist from cracking. The process was repeated with mask (3) to create a second 4-μm layer, with diluted SU8-2005 (90% v/v in cyclopentanone). After fabrication, the mold was treated once with silane Trichloro(1H,1H,2H,2H-perfluorooctyl)silane (448931, Sigma-Aldrich) to prevent the PDMS from sticking and subsequently damaging the constrictions. A mold is typically used at least ten times. A 10:1 mixture of PDMS and cross-linking agent (Sylgard 184, Dow Corning) was poured on the master mold. After degassing in a vacuum chamber for 30 min and curing at 80°C from 2 h up to overnight, the PDMS microchannel was peeled off from the mold.

#### Bottom coverslip

Glass coverslips were cleaned by sonication in acetone then in isopropanol, for 30 min each, washed in ethanol for 5 min, and rinsed abundantly with MilliQ water. Cleaned coverslips were stored in MilliQ water before use. A few drops of PDMS were poured onto a dried coverslip and spin-coated at 4000 rpm for 45 s before being cured at 80°C overnight.

#### Chip assembly

The three inlets and the outlet of the PDMS microchannel were drilled with a 0.75-mm punch (Biopsy Punch 504529, World Precision Instruments) prior to assemble the microfluidic chip by bonding the microchannel to the bottom coverslip via plasma treatment, followed by a curing step at 80°C overnight.

### Microfluidic experiments

The experiments were performed on a microscope (IX71, Olympus) equipped with a 20x objective and high-speed cameras (Fastcam Mini UX, Photron and VEO310, Phantom) at 37°C. Prior to experiments, cells were cultured for 72 h (including 48 h of Atazanavir treatment), to ensure ≈90% cell confluence. The microfluidic chip inlets and oulet were connected via Teflon tubing (0.012” inner diameter (ID) and 0.030” outer diameter (OD), BB311-30, Scientific Commodities Inc.) to a pressure controller (MFCS™-EZ 1000 mbar, FLUIGENT) that allows to fix the pressure drop between the inlets and outlet and thus the circulation of cells/nuclei in suspension. Microchannels were passivated with 10% Pluronic®F-127 (P2443, Sigma) in PBS for 1 h at room temperature. Inlets (1) and (3) were used to inject buffer solutions while cells/nuclei were injected with inlet (2) (see Fig. 2A for a schematic view of the chip).

#### Cell experiments

Cells suspended in complete medium supplemented with 10 mM HEPES, 1% BSA (± cytoskeleton drugs) were injected in microchannels under a pressure drop of ΔP = 165 mbar (P_inlets_ = 170 mbar, P_oulet_ = 5 mbar). Cells were observed in brightfield and movies were acquired at frame rates varying from 125 to 1000 fps.

#### Isolated nucleus experiments

Isolated nuclei suspended in PBS with 1% BSA were injected into microchannels under a pressure drop of ΔP = 200 mbar (P_inlet_s = 205 mbar, P_oulet_ = 5 mbar). Nuclei were observed in epifluorescence and movies were acquired between 500 and 1000 fps.

### Image processing and analysis

#### Cell analysis

Movies of cells were pre-processed using FIJI software to enhance contrast and segment the cells. To enhance cell contrast and remove constrictions (static in the movie), a median image from a few frames was computed and a new movie was produced by subtracting the median image. Using Non-Local Mean Denoising Plugin (Darbon et al., 2008; Buades et al., 2011), the background noise was removed while keeping the cell intact, which was then manually thresholded. Using Matlab software (MathWorks, r2018b version), a bounding box was fitted around the thresholded cell at each time point to measure the cell dimensions (length L along the flow axis and width w perpendicularly to it) over time.

#### Isolated nucleus analysis

Movies of nuclei were pre-processed using FIJI software, with automated tracking of individual nuclei using Trackmate (Jaqaman et al., 2008). Each track contained the X-Y positions of a nucleus centroid at each timepoint. Using the X-Y positions of the centroids, we defined an initial mask of 12×12 pixel^2^ that evolved to perfectly fit the nucleus contour using the Active Contour method in Matlab (Lankton and Tannenbaum, 2008; Getreuer, 2012). Once the final contour was retrieved, the image was segmented and the bounding box analysis was performed such as for cells.

#### Cellular and nuclear volumes

Additionally, volumes of cells and isolated nuclei were computed from microfluidic images before contact with constrictions, assuming they have a “pancake” shape. The volume is computed as the projected area multiplied by the microfluidic channel height: V = area × h_1_.

### Tongue length analysis

#### Entry time and slope analysis

With a home-made Matlab routine, the time evolution of the tongue length l(t) was plotted and four parameters were manually determined: the entry time T_e_ (in seconds), defined as the time to fully enter a constriction, and three slopes S_I_ to S_III_ (in s^− 1^) corresponding to three distinct regimes displayed during the cell entry and considered as linear in a first approximation. To calculate the slopes, the first and last points of each regime were selected, and a tangent was automatically fitted that provided the slope value.

#### Rheological fit

Matlab fit function (MATLAB 2018) was used on the curves l(t), with expression a(1-exp(-bt)) + ct, where a, b, and c are fit parameters; the combination of these parameters provides the physical quantities defined in Eq. (8). We performed a series of 6 fits of the experimental curves with a set of time intervals I_m_ = [0;m × T_e_/10] with m = 4, …, 10, where T_e_ is the cell entry time. For each fit, we evaluated the corresponding physical parameters and R2 chi-square values, and defined for each cell the following lists: (η2^(m)^)_m= 4,…,10_, (η1^(m)^)_m= 4,…,10_, (E^(m)^)_m=4,…,10_ and (R^2(m)^)_m=4,…,10_. We only retained fitted parameters whose corresponding fits had the maximal R^2^ value that is larger than a critical threshold set at 0.85, i.e. those such that R^2^_I_ = max(R^2^_m_)_m=4,…,10_ > 0.85. We discarded too large values of η_2_ > 10^5^ Pa.s, η_1_ > 500 Pa.s and E > 70 kPa, threshold values one to two orders of magnitude larger than typical median values. We also discarded too low values of η_1_ < 5 Pa.s and E < 0.7 kPa.

### Rheological models

Here we show how we derive the elastic modulus and the viscosities of cells using a viscoelastic model. We follow a method laid out in previous works using micropipettes (Guevorkian et al., 2010; Davidson et al., 2019), with adaptation for a squared cross-section constriction.

#### Constant pressure drop approximation

In Guevorkian et al. (Guevorkian et al., 2010), aspiration is performed using a cylindrical pipette of circular cross-section 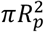 Considering volume conservation (experimentally, the cell volume decreases at most by 20%), the aspiration force under a pressure drop ΔP between the exterior and the interior of the pipette reads as:

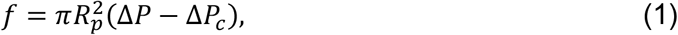

where ΔP_c_ is the Laplace pressure, i.e. the critical pressure required to push cells through constrictions, which reads as:

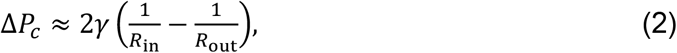

where the cell radii R_in_ = R_p_ and R_out_ correspond to the portion of the cell either inside (front of the cell) or outside (back of the cell) the constriction, respectively, and γ is the cortical surface tension.

Here we argue that the Laplace pressure ΔP_c_ can be neglected. Indeed, the cortical tension lies in the 5.10^−4^ N.m^-1^ (cortical tension in fibroblast cells (Tinevez et al., 2009; Salbreux et al., 2012)), with upper values reported at 2.10^−3^ N.m^-1^ (dividing Hela cells (Fischer-Friedrich et al., 2014)), corresponding to ΔP_c_ in the 0.5 to 2 kPa range. In comparison, the applied pressure drop is ΔP = 165 mbar = 16.5 kPa, which is significantly larger than ΔP_c_. Considering a more realistic geometry with a Laplace pressure given as in Bruus et al. (Bruus, 2008) does not affect our conclusion. In addition, the validity of our approximation is verified a posteriori through the estimation of a capillary number based on the measurement of the cell viscosity. We point out that Dupire et al. (Dupire et al., 2020), who focused on the measurement of the surface tension, considered a significantly lower applied pressure. Therefore, in the rest of the manuscript we consider that:

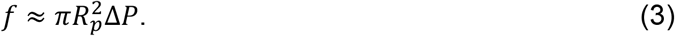

#### Jeffreys rheological model

The established literature (Guevorkian et al., 2010; Davidson et al., 2019; Wintner et al., 2020) models the evolution of the deformation of various objects (from nuclei to cell aggregates) in micropipettes through the Jeffreys rheological model. The Jeffreys model results in the following time evolution of the aspirated tongue length (Davidson et al., 2019):

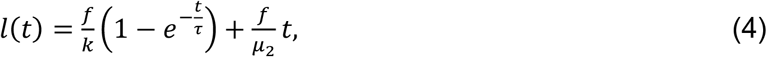

where *f* is the applied force, defined in Eq. (1), k is a spring constant, τ is a relaxation time and μ_2_ is the long-time viscosity. The first term in Eq. (4) describes the short time behavior, during which the pressure force is balanced by the cell elastic forces. The cell yields after a characteristic time τ, which we can express in terms of a short-time viscosity μ_1_ as τ = k/μ_1_. The second term in Eq. (4) describes the long-time behavior (before complete cell entrance is achieved) during which the pressure force is balanced by the viscous force due to a plug flow at the entrance of the capillary, leading to a linear increase of the regime strain with time. The stress/strain relationship of Eq. (4) is commonly represented diagrammatically in terms of a Kelvin-Voigt element, i.e., a spring (k) and a dashpot (μ_1_ = k/τ) in parallel, in series with a dashpot (μ_2_).

#### Relation to rheology

Here we propose to map the spring k, viscoelastic time τ and viscosity μ_2_ parameters defined in Eq. (4) to a simplified model for the cell rheological properties. Following the literature (Guevorkian et al., 2010; Davidson et al., 2019), we discuss the expected behaviour of the cell considered as a homogeneous viscoelastic material under aspiration through a cylindrical constriction of radius R_p_.

At short time scales, the aspiration force is balanced by the elastic deformation and is given by the following equation:

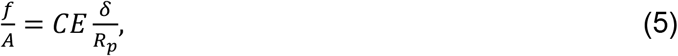

where A = π R_p_2 is the cross-sectional area of the constriction, C is a geometrical factor approximately equivalent to 1 (Guevorkian et al., 2010; Davidson et al., 2019), E is the elastic modulus and δ is the elastic deformation at short times. We thus obtain *f* = π R_p_ E δ. The spring constant k is the coefficient between the applied force f and the extension δ, f = k·δ, leading to the relation:

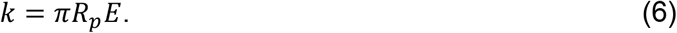

At long time scales, the aspiration force is balanced by two dissipation forces. The first one arises from the viscous flow of the cell through the entrance of the constriction, such as the flow of a viscous fluid through a circular pore described by Sampson (Sampson, 1891). The second source of dissipation arises from the resistance of the tongue slipping along the constriction wall. Combining both terms leads to (Guevorkian et al., 2010; Piroird et al., 2011):

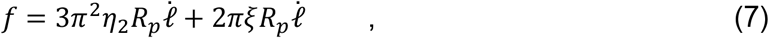

where η_2_ is the cell viscosity and ξ is the cell-wall friction coefficient. As in Refs. (Guevorkian et al., 2010) and (Davidson et al., 2019), we can ignore the friction on the surface of the constriction: indeed, once the cell has fully entered into the constriction, its velocity (in the 50 µm.s^-1^ range) is significantly larger than during the deformation phase. Identifying 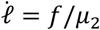 from the second term in Eq. (4) leads to:

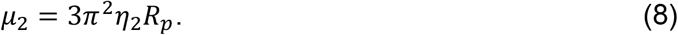

Inserting Eqs. (6) and (8) into Eq. (4), we obtain the following relation of the cell tongue length as a function of the cell rheological properties:

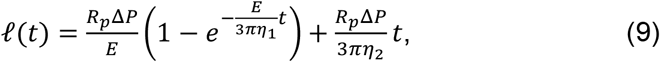

which is the expression that we use within the main text.

A key interest of Eq. (9) is to provide a set of intensive mechanical properties that allows comparison between classes of cell types. For each cell type, we verified that the estimated mechanical parameters in Eq. (9) did not vary for cells of different sizes.

#### Effective radius of the constriction

For a cylindrical pipette as in Guevorkian et al. (Guevorkian et al., 2010), the pipette radius R_p_ is used in Eq. (1). In our case, the constrictions are rectangular instead of cylindrical and R_p_ is replaced by an effective radius R_eff_ that is the function of the height and width of the rectangular cross-section. Following Davidson et al. (Davidson et al., 2019) and the theoretical study from Son (Son, 2007), we express the effective radius as:

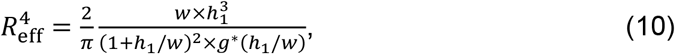

where h_1_ and w are the dimensions of the constriction cross-section and g* is a dimensionless function of the form:

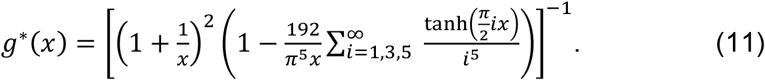

With w = h_1_ = 6 μm, applying Eq. (10) yields R_eff_ = 4.32 μm.

We point out that Eq. (10) is not strictly exact in our case, since our geometry is off-centered in height, which is not exactly the case considered by Son (Son, 2007). Nonetheless, we expect R_eff_ to be a satisfactory approximation. In addition, in our study the constriction geometry is identical for all cell types and drug treatment; any approximation on the value of R_eff_ will not affect conclusions based on the comparisons of mechanical parameters between experiments.

#### A simple power law between cell initial length L_0_ and entry time T_e_ allows for crossed comparisons of viscous moduli

Here we propose to extract a simple relation between the entry time T_e_ and the cell radius before entry that holds for sufficiently large cells. In this case T_e_ ≫ τ on average and, by neglecting the time spent in both phases I and III (see Fig. 2C), the aspirated tongue length at time T_e_ derives from Eq. (9) as:

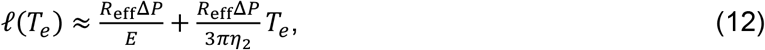

Assuming the cell has a disk shape of diameter L_0_ before entering the constriction, the initial volume V_i_ can be expressed as V_i_ = π (L_0_/2)^2^ h_1_. Once fully entered in the constriction, the cell shape is approximately a parallelepiped of final length l(T_e_) and final volume V_f_ ≈ l(T_e_) w h_1_. Considering the cell volume to be conserved, the final aspirated tongue length l(T_e_) can be expressed in terms of L_0_ as:

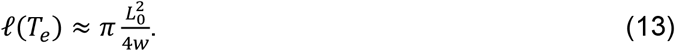

Substituting the latter expression into Eq. (12), we obtain, for large cells L_0_/w ≫ 1, the following approximate scaling of the entry time T_e_ with the initial cell diameter (or initial length) L_0_:

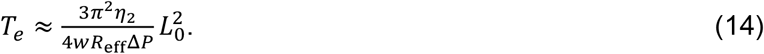

Eq. (14) provides a rationale for fitting the relation between the experimental entry time T_e_ and the cell diameter L_0_ through the following log-log scaling:

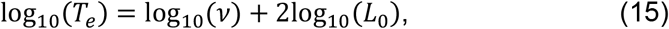

where we expect the quantity ν to be proportional to the long-time cell viscosity η_2_ (see Fig. 10B). We measure the value of ν for all cells whose experiment time is long enough (T_e_ > 10τ ≈ 0.3 s).

#### Alternative model: power-law rheology

Following Refs. (Davidson et al., 2019; Dahl et al., 2006), we also evaluate the efficiency of a power-law fit of the tongue length data. We employ a similar fitting procedure to the one previously used for viscosity extraction, with a master curve:

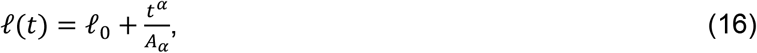

where *l*_0_ corresponds to the very short-time elastic response and A_α_ is a quantity that corresponds to a non-Newtonian viscous coefficient (in the limit α = 1, A_α_ corresponds to a Newtonian viscosity). Inspired by the range of values measured in Ref. (Davidson et al., 2019), we set the value of the exponent to α?= 0.5. We then observe that the relations between the 1/A_α_ prefactors in all cell types also correspond to the previously measured relations on η_2_ (see Fig. S8).

## Statistical analysis

When indicated, N, n, and n’ are the number of experiments, of analyzed cells, and of fitted curves, respectively. Statistical calculations were performed using Prism 6.0 statistical software (GraphPad). Significant differences (p-values) are indicated with *(or #,¤): *, p < 0.05; **, p < 0.01; ***, p < 0.001 ; ****, p < 0.0001.

For experiments examining the proportions of cells with nuclear abnormalities or SA-β-galactosidase positive staining, data were averaged over at least 3 independent experiments (mean ± SD, standard deviation) and mean values were compared using the Fisher’s exact test and the Bonferroni post-hoc analysis to correct for multiple comparisons.

For BrdU incorporation, data were averaged over at least 3 independent experiments (mean ± SD) for each condition, and mean values were compared using the Kruskal-Wallis test followed by the Dunn’s multiple comparison post-hoc analysis when appropriate.

For nuclear and cellular volumes, the median values and 95% Confidence Intervals (CIs) were calculated from measurements on individual cells pooled from at least 2 independent experiments, and median values were compared using the Kruskal-Wallis test followed by the Dunn’s multiple comparison analysis.

Entry time, tongue length slopes, and rheological parameters derived from the fits of the tongue length curves all displayed non-Gaussian distributions. The median values were calculated with 95% CIs from individual cells pooled from at least 3 independent experiments, and data were compared using the Kruskal-Wallis test followed by the Dunn’s multiple comparison analysis.

## Online supplemental material

Table S1 displays the rheological parameters measured upon actin and microtubule networks disruption.

Table S2 presents a summary of fibroblast cell lines used in the study.

Figure S1 shows the cellular and nuclear volumes of cells measured from confocal imaging.

Figure S2 shows the dot plots of cell entry times in constrictions, unsorted (corresponding to Fig. 3A) or sorted by cell size.

Figure S3 shows the slopes S_I_ to S_III_ of the 3 entry phases approximated as linear.

Figure S4 shows the dot plots of the rheological parameters, corresponding to the bar plots in Fig. 3D-F.

Figure S5 shows the similar behavior of cell lines obtained either with the complete Jeffreys model or the simplified Jeffreys model.

Figure S6 shows fits of cell tongue length using the power law model.

Figure S7 shows cells at different time points of entry in constrictions, with blebs at the front. Figure S8 shows the dot plots of nucleus entry times sorted by nucleus size.

Figure S9 shows epifluorescence images of cells upon actin, microtubule, and both actin and microtubule networks disruption.

Figure S10 schematizes the workflow of the master fabrication for molding the microfluidic chip.

Video S1 shows an FPLD2 cell entering a constriction simultaneously with the elongation of the cell tongue.

Video S2 shows typical movies of an untreated control cell and an Atazanavir-treated cell entering constrictions.

Video S3 shows the entry of an isolated nucleus in a constriction.

## Acknowledgments

The project leading to this publication has received funding from Excellence Initiative of Aix-Marseille University - A*MIDEX (A-M-AAP-ID-17-66-170301-11.30) and from France 2030, the French Government program managed by the French National Research Agency (ANR-16-CONV-0001). Corinne Vigouroux is a member of the European Reference Network on Rare Endocrine Conditions – Project ID No 739527.

## Author contributions

EH, AV and CB designed the study. CJ, AAV and LHC performed the experiments. AL, FB and CD contributed to setting up the experiments. CV and MCV provided human cell lines. CJ, AAV, and MK analyzed data. CJ, AAV, MK, AV, CB, JFR, and EH interpreted data. CJ, AAV, MK, JFR, CB and EH wrote the paper. All authors have revised the manuscript and agreed to the final version.

The authors declare no competing financial interests.

## Notes

### Competing Interest Statement

The authors have declared no competing interest.

### Summary of Updates

Manuscript revised for clarification

