## Supplementary Tables and Figures for "Enhanced cell viscosity: a new phenotype associated with lamin A/C alterations"

### Supplementary information for: Enhanced cell viscosity: a new phenotype associated with lamin A/C alterations

**This PDF file includes:**
Tables S1 to S2
Figures S1 to S10
Legends of Videos S1 to S3

| UNT | | | | |
|---|---|---|---|---|
| | DMSO | N | L | NL |
| $T_e$ (s) | 0.48 [0.34-0.60] | 0.26 [0.21-0.33] | 0.22 [0.18-0.31] | 0.15 [0.11-0.18] |
| $\eta_1$ (Pa.s) | 26 [22-29] | 29 [24-32] | 9 [7-11] | 10 [8-12] |
| $E$ (kPa) | 5.7 [5.1-6.1] | 6.2 [5.6-6.6] | 4.5 [4.2-4.9] | 4.5 [4.0-5.0] |
| $\eta_2$ (Pa.s) | 251 [217-318] | 287 [242-335] | 146 [112-191] | 91 [78-116] |
| AZN | | | | |
| | DMSO | N | L | NL |
| $T_e$ (s) | 1.75 [1.33-2.12] | 0.95 [0.75-1.20] | 0.49 [0.42-0.66] | 0.43 [0.36-0.55] |
| $\eta_1$ (Pa.s) | 25 [23-28] | 34 [28-39] | 10 [8-11] | 13 [6-16] |
| $E$ (kPa) | 4.0 [3.9-4.3] | 4.7 [4.4-5.0] | 4.1 [3.8-4.3] | 3.9 [3.6-4.3] |
| $\eta_2$ (Pa.s) | 700 [582-844] | 416 [333-506] | 244 [190-302] | 181 [138-227] |
| K | | | | |
| | DMSO | N | L | NL |
| $T_e$ (s) | 1.26 [0.96-1.49] | 0.56 [0.42-0.76] | 0.54 [0.40-0.68] | 0.41 [0.29-0.59] |
| $\eta_1$ (Pa.s) | 23 [19-27] | 23 [19-25] | 11 [8-13] | 11 [9-14] |
| $E$ (kPa) | 4.1 [3.7-4.5] | 4.2 3.8-4.6] | 3.8 [3.6-4.4] | 3.9 [3.4-4.3] |
| $\eta_2$ (Pa.s) | 631 [541-725] | 293 [247-343] | 219 [171-304] | 168 [142-219] |

**Table 1. Changes in rheological parameters upon actin and microtubule networks disruption in UNT, AZN and K cells (medians [95% CI]).** Colors indicate significant differences between N/L/NL *vs* DMSO conditions: ns * ** *** ****. N = nocodazole, L = latrunculine A, NL = nocodazole + latrunculin A.

| Cell | Denotation | Description | Age | Sex | Biopsy source | Origin |
|---|---|---|---|---|---|---|
| AG07095 | UNT | Apparently healthy individual | 2 | M | Foreskin | Purchased from Coriell Institute |
| AG06917 | Progeria | *LMNA* G608G mutation, Progeria | 3 | M | Arm | Purchased from Coriell Institute |
| Patient M* | FPLD2 M or M | *LMNA* R482W mutation, FPLD2 | 36 | F | Skin | patient 12 (1) |
| Patient K* | FPLD2 K or K | *LMNA* R482W mutation, FPLD2 | 41 | F | Skin | patient 8 (1) |
| Patient T2D* | T2D | No *LMNA* mutation, diabetic | 49 | M | Skin | patient 17 (2) |

**Table 2. Summary of fibroblast cell lines.**

* Patient 8 and 12 cells from Ref. (1), patient 17 cells from Ref. (2). Informed consent was obtained from all patients. Patients were unrelated.

Patients K and M, affected by FPLD2 associated with R482W mutation in the *LMNA* gene, presented similar symptoms such as peripheral lipoatrophy, fat accumulation in face and neck, muscular hypertrophy, fatty liver, hypertriglyceridemia, diabetes. Patient K had a more severe form of Dunnigan syndrome than patient M: lower amount of body fat, antecedent of acute pancreatitis at age 19, very severe insulin resistance, and diabetes was complicated by retinopathy and nephropathy. Patient T2D, who does not carry any *LMNA* mutation, presented neuromuscular complaint, fatty liver, severe hypertriglyceridemia, and diabetes.

1. C Vatier, et al., One-year metreleptin improves insulin secretion in patients with diabetes linked to genetic lipodystrophic syndromes. *Diabetes, Obes. Metab.* **18**, 693–697 (2016).

2. A Dutour, et al., High prevalence of laminopathies among patients with metabolic syndrome. *Hum. molecular genetics* **20**, 3779–3786 (2011).

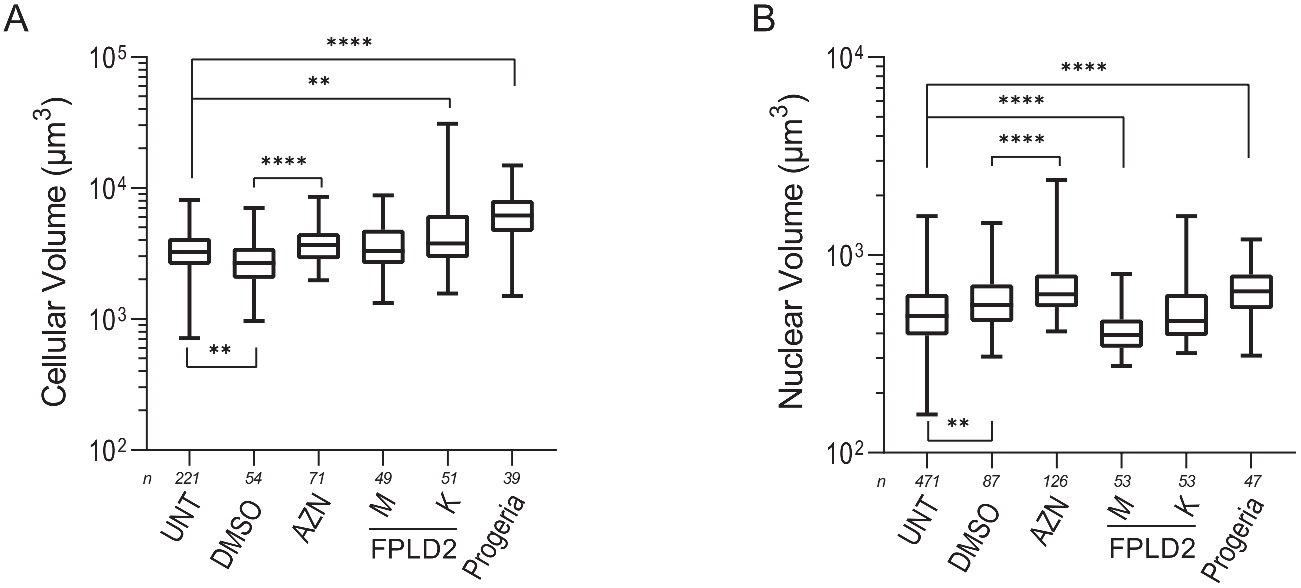


C

| Cell | Cellular Volume ($\mu$m$^3$) | | Nuclear Volume ($\mu$m$^3$) | |
|---|---|---|---|---|
| | Median | 95% CI | Median | 95% CI |
| UNT | 3254 | 3068-3457 | 491 | 469-511 |
| DMSO | 2678 | 2294-3022 | 558 | 511-598 |
| AZN | 3677 | 3192-4205 | 633 | 591-686 |
| FPLD2 M | 3310 | 2986-3731 | 393 | 350-432 |
| FPLD2 K | 3775 | 3490-4495 | 462 | 413-578 |
| Progeria | 6171 | 4976-7635 | 654 | 568-737 |

**Fig. 1. Cellular and nuclear volumes of control and altered lamin A/C cells.** UNT: untreated control cells; DMSO: control cells incubated with DMSO for 48 h; AZN: control cells treated with Atazanavir (in DMSO) for 48 h; M,K: cells from two patients with FLPD2 carrying the lamin A/C R482W mutation; Progeria: cells from a patient with HGPS carrying the lamin A/C G608G mutation. Adhered cells were labelled for plasma membrane (with Cell Brite) and nuclei (with Hoechst) before detachment and resuspension in culture medium. Cellular and nuclear volumes were computed from confocal imaging. A-B) Boxplot representations of the cellular (A) and nuclear (B) volumes, with median values, 25% and 75% percentiles, and min/max values as whiskers. Number of experiments: N $\geq$ 2; number of analyzed cells: *n*. C) Table recapitulating median values of the volumes and 95% Confidence Intervals (CIs).

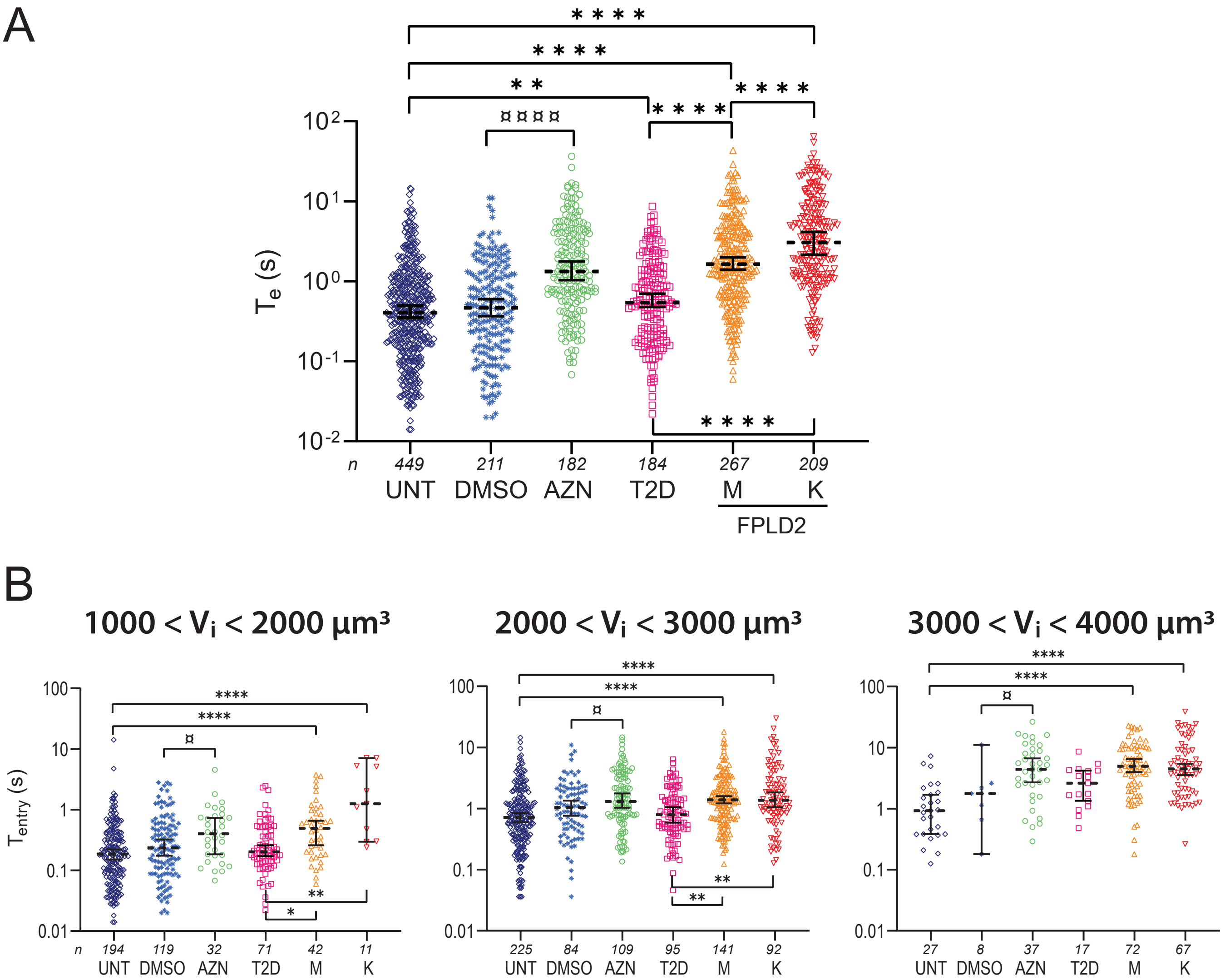


**Fig. 2. Entry time in constrictions $T_e$ is larger in altered lamin A/C cells independently of cell size.** Cell types as in Fig. 3: UNT: untreated control cells; DMSO: control cells incubated with DMSO for 48 h; AZN: control cells treated with Atazanavir for 48 h; T2D: cells from the diabetic patient; M,K: cells from the patients with FPLD2. A) Dotplot representation of $T_e$, independently of cell size. The non-Gaussian distributions are plotted in log scale; median values and 95% CIs are indicated. The plots correspond to the bar plots in Fig. 3A and illustrate the number of analysed cells. B-E) Same plots as in (A), with cells sorted into three size populations. The cellular volumes were computed from the images of the cells before they enter the microfluidic constrictions. Both types of cells with lamin A/C alterations, FPLD2 and AZN, display larger $T_e$ compared to their respective controls, whatever the volume range. Significant differences: FPLD2 and L *vs* UNT cells, $^*$; AZN *vs* DMSO cells, ¤. Number of experiments: N $\geq$ 3; number of analyzed cells: *n*.

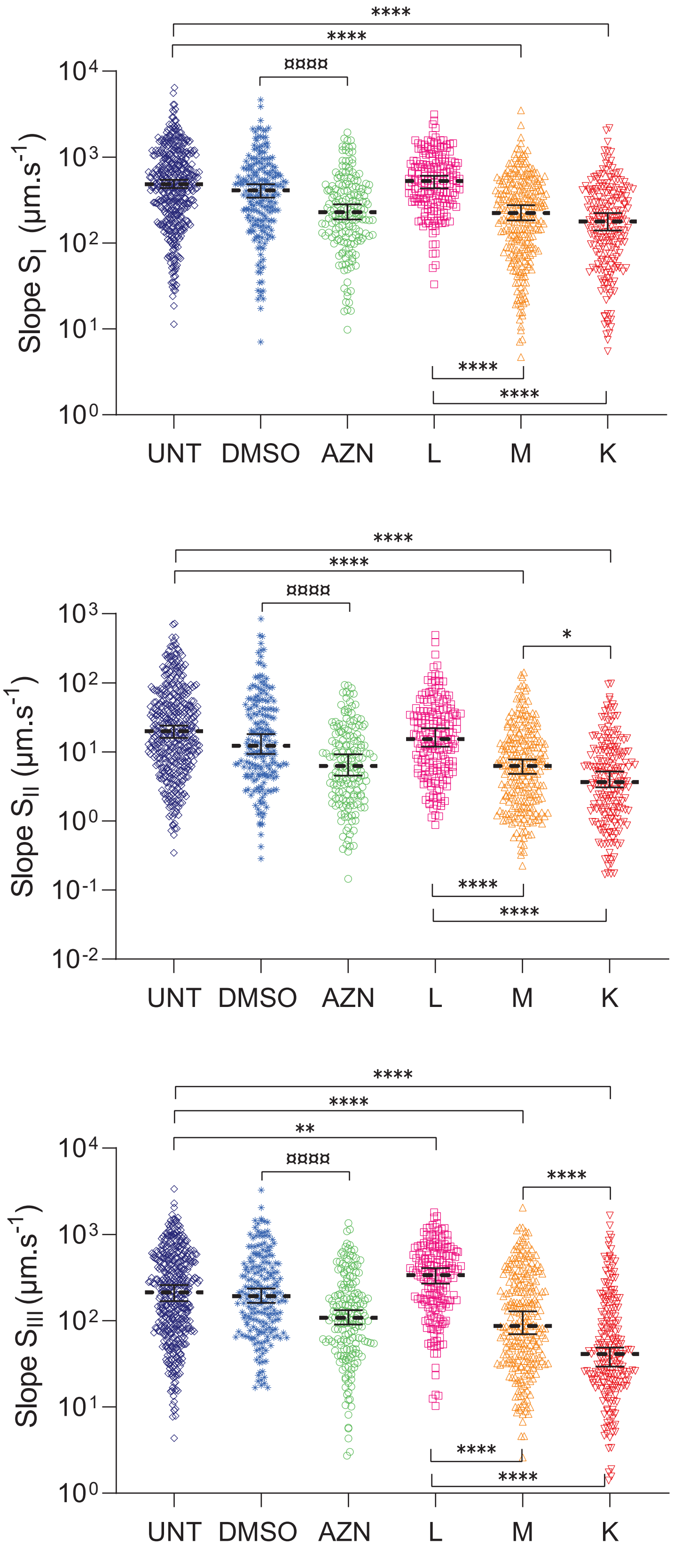


**Fig. 3. The linear slopes $S_I - S_{III}$ of the three regimes in the temporal curve of the tongue length $\ell(t)$ are changed by the cell condition, consistently with the entry time $T_e$.** Cells as in Figs. 3 and S2. Significant differences: FPLD2 and T2D *vs* UNT cells, *; AZN *vs* DMSO cells, ¤. Number of experiments: N $\geq$ 3; number of analyzed cells: *n* > 150.

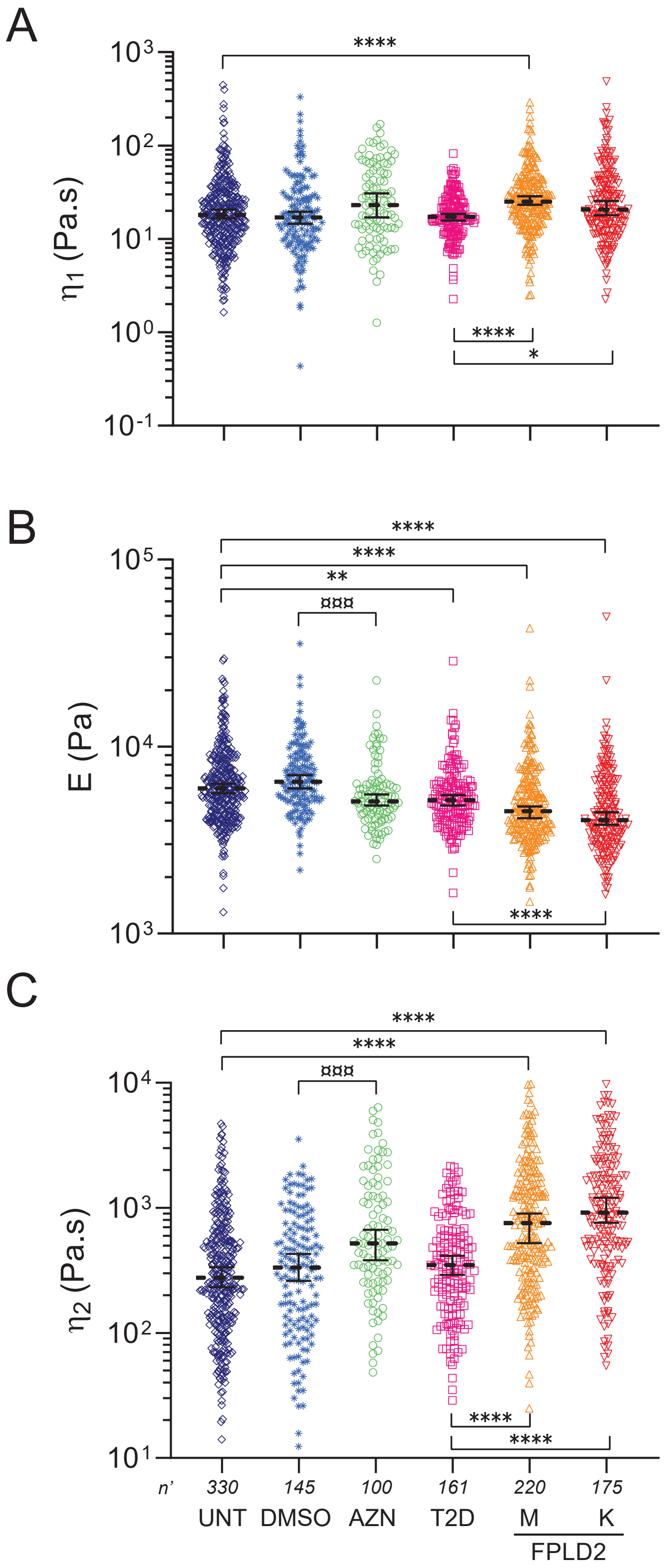


**Fig. 4. Dotplot representations of cell viscoelastic parameters extracted with the rheological model.** Cells as in Figs. 3, S2 and S3. The plots correspond to the bar plots in Fig. 3D-F and illustrate the number of analyzed cells. A) Short-time viscosity $\eta_1$. B) Elastic modulus $E$. C) Long-time viscosity $\eta_2$. Number of experiments : N $\geq$ 3 ; number of fitted curves: *n'*.

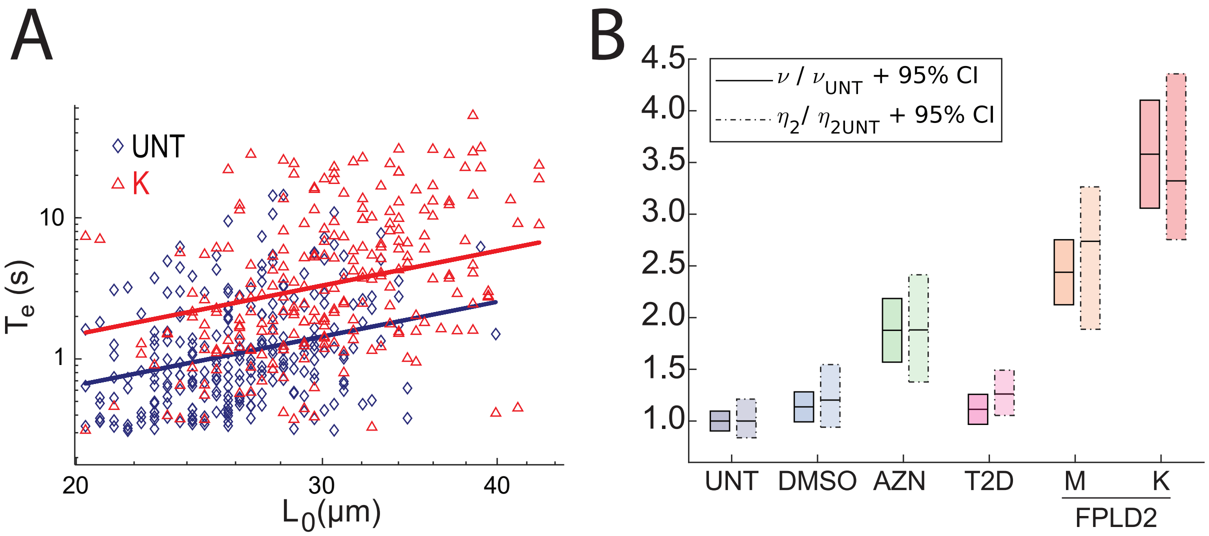


**Fig. 5. Alteration of long time viscosity is detected by a simple relationship between entry time and cell initial length.** (A) Dependence of the entry time $T_e$ with the cell initial length $L_0$ for UNT and K cells. Data are plotted in log-log scale and fitted with the scaling law $T_e \sim \nu L_0^2$. Cells with $T_e$ < 0.3 s are not considered. (B) Boxplots of the constant $\nu$ extracted from the scaling law fit (solid line) and of long-time viscosity $\eta_2$ (dashed line), normalized to the values obtained for UNT cells (median values $\pm$ 95% CIs).

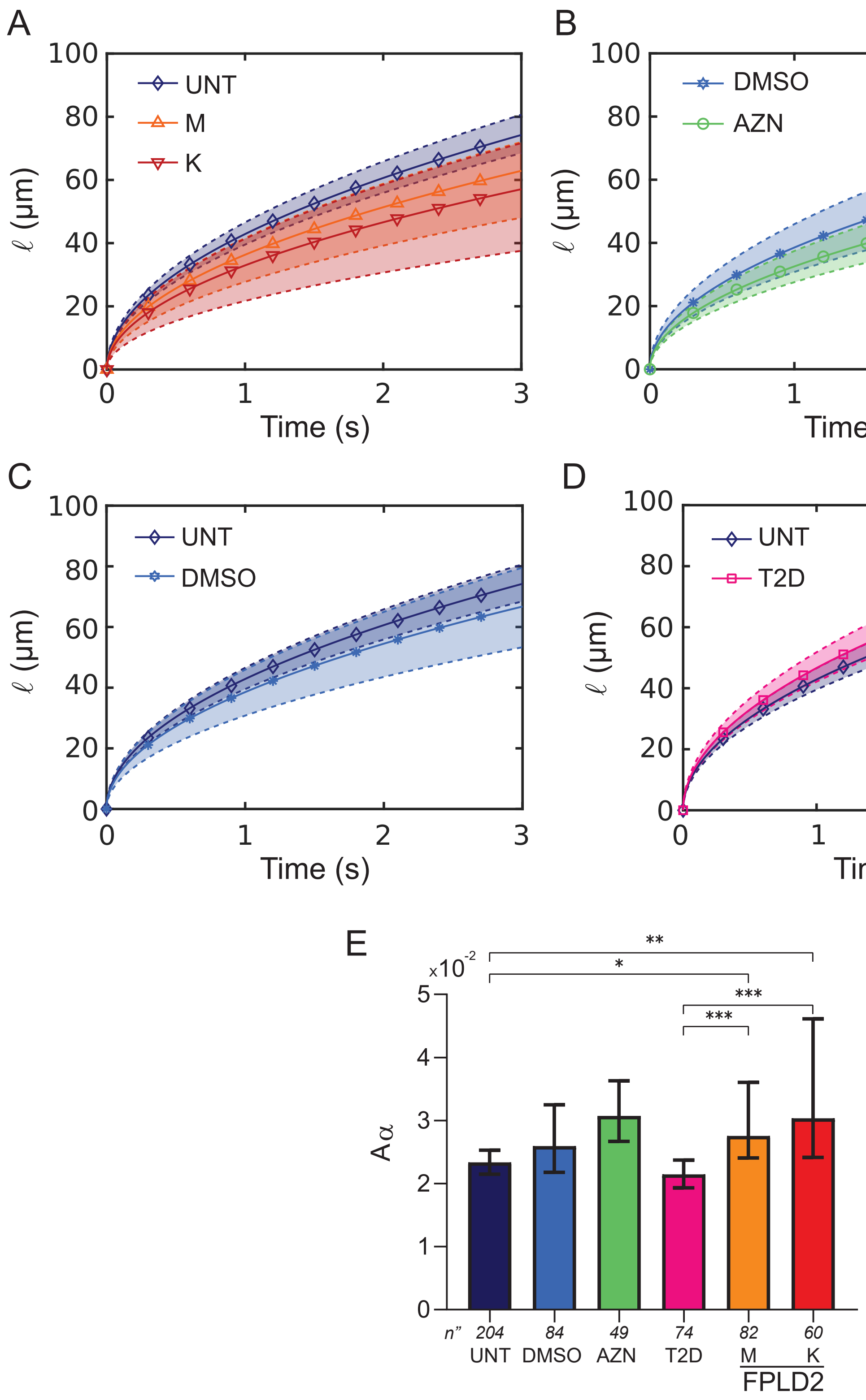


**Fig. 6. Tongue length fits using a power law.** (A-D) Power law fits $\ell = \ell_0 + t^{0.5}/A_\alpha$ for (A) UNT *vs* M and K cells, (B) DMSO *vs* AZN cells, (C) UNT *vs* DMSO cells, and (D) UNT *vs* L cells. Solid curves and symbols indicate the median fits plotted with the median values of the parameters, dashed upper and lower curves delineate the 95% CIs. (E) Comparison of $A_\alpha$ values extracted from the power law fits for the different cell types (medians $\pm$ 95% CIs). Number of experiments: N $\geq$ 3; number of fitted curves: *n”*.

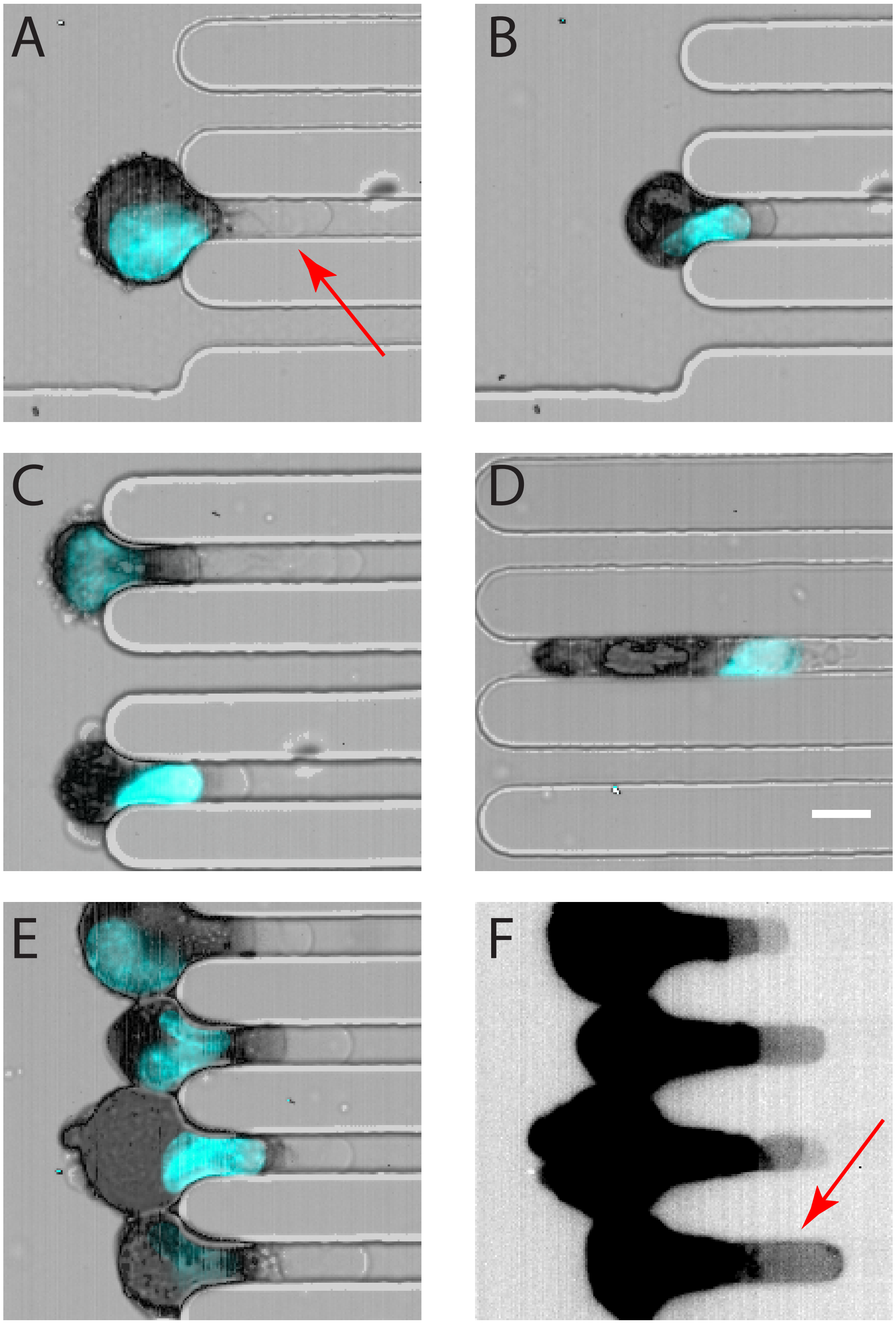


**Fig. 7. The cell forms a bleb at the front that precedes the actin cortex and the nucleus during deformation in the constriction.** The cell membrane, actin and nucleus are imaged at different timepoints during entry in constriction. A) The cell comes into contact with the constriction and a bleb immediately enters (phase I). B) The bleb moves forward, the actin cortex and the nucleus slowly deform (phase II). C) The nucleus has completely entered the constriction and the rear of the cell deforms (bottom cell), the bleb is still present at the front (phase III). D) The cell is fully deformed and transits through the constriction, with the bleb at the front followed by actin, nucleus and cell rear (phase IV). E-F) Saturating the image highlights the transparent front bleb, with the actin in black. Cell membrane is imaged in brightfield; nucleus and F-actin are imaged in epifluorescence, via Hoechst (blue) and SPY-actin 555 (black) labelling, respectively. Red arrows indicate some blebs. Scale bar = 10 $\mu$m.

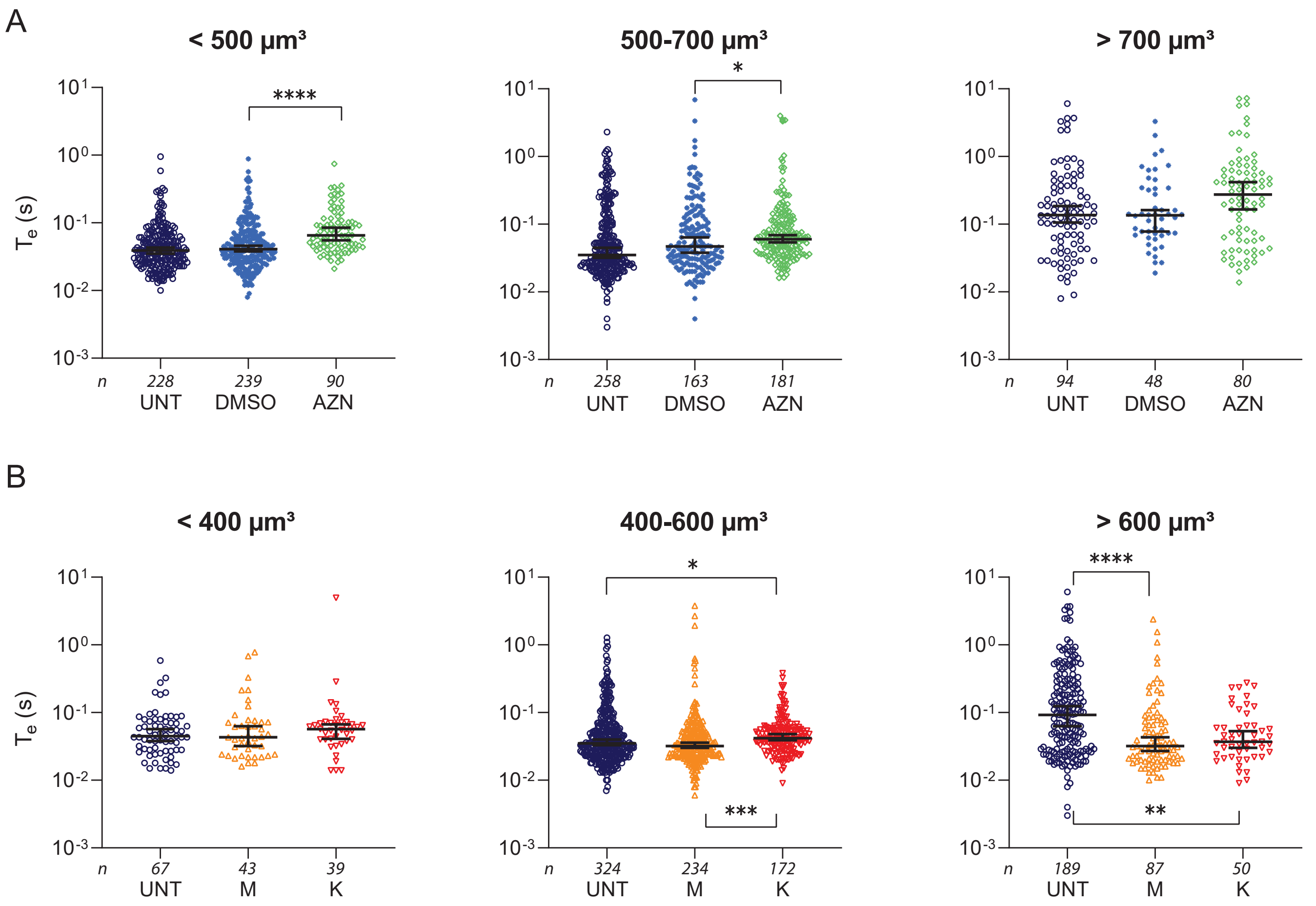


**Fig. 8. Entry time of isolated nuclei in constrictions sorted by nucleus volume.** Dotplot representations of entry times $T_e$ sorted in three size populations that were defined based on the median volumes of nuclei from AZN cells ($\approx$600 $\mu$m$^3$, B) and from FPLD2 M/K cells ($\approx$500 $\mu$m$^3$, C). Number of experiments: N $\geq$ 4; number of analyzed nuclei: *n*.

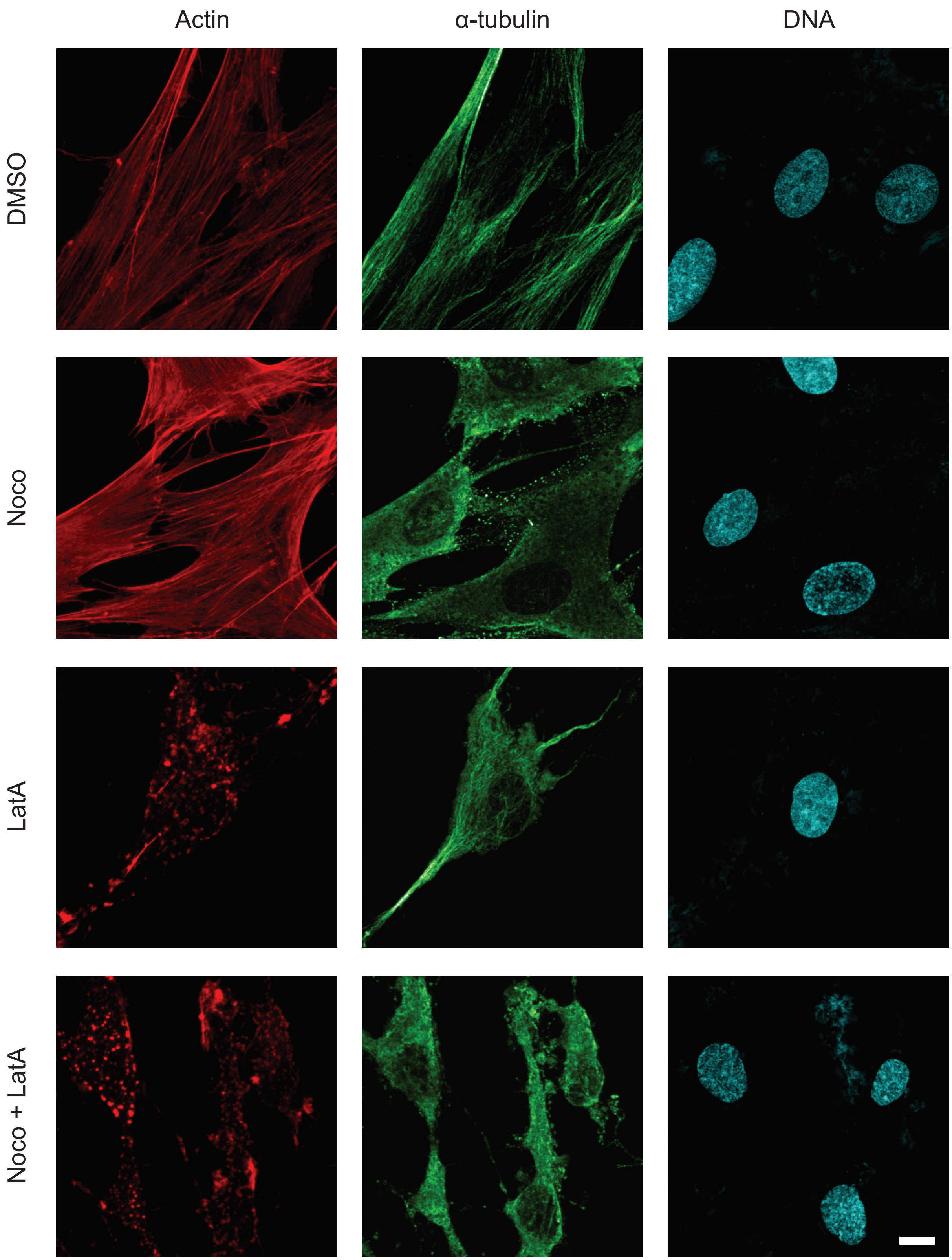


**Fig. 9. Actin and microtubule network depolymerization upon cytoskeletal drug treaments.** Maximal intensity projection from confocal z-stacks of F-actin (red), $\alpha$-tubulin (green) and DNA (blue). Adhered UNT cells were incubated with DMSO, or treated with nocodazole (Noco), latrunculine A (LatA) or both nocodazole and latrunculine A (Noco+Lat A) prior to immunostaining. Scale bar: 10 $\mu$m.

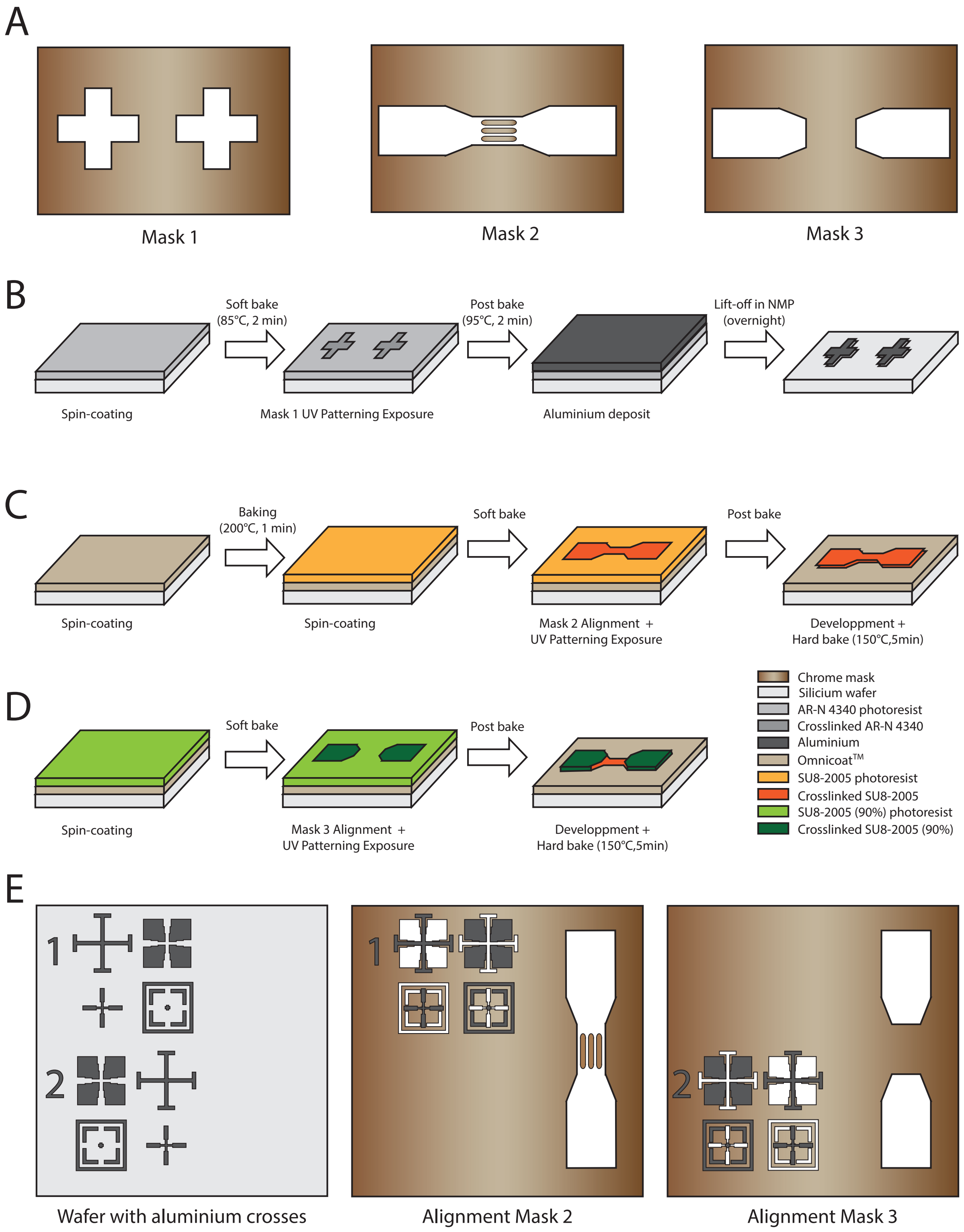


**Fig. 10. Workflow of master mold fabrication.** A) Simplified schematics of the three masks used to create the mold: Mask 1, alignment crosses; Mask 2, main channel with micrometer-sized constrictions in the central part; and Mask 3, main channel without constrictions. B-D) Steps to create the two-height master mold. E) Alignment of masks 2 and 3 prior to UV patterning exposure.

**Video legends**

**Video S1.** Entry of a 24-$\mu$m FPLD2 K cell and corresponding time evolution of the tongue length.

**Video S2.** Entry and transit of fibroblasts in 6x6 $\mu m^2$ constrictions: top) 24-$\mu$m untreated control cell, middle) 23-$\mu$m Atazanavir-treated cell, bottom) 27-$\mu$m FPLD2 K cell. Cells were observed in brightfield microscopy, the background with the image of the constrictions has been subtracted to enhance cell contrast. Time is indicated in sec. Scale bar: 10 $\mu$m.

**Video S3.** Entry and transit of a 20-$\mu$m nucleus in a 3x3 $\mu m^2$ constriction. Nucleus is labelled with Hoechst and observed in epifluorescence microscopy. Time is indicated in msec. Scale bar: 10 $\mu$m.
